# Convergent and divergent responses of the rhizosphere chemistry and bacterial communities to a stress gradient in the Atacama Desert

**DOI:** 10.1101/2023.10.16.562209

**Authors:** Thomas Dussarrat, Claudio Latorre, Millena C. Barros Santos, Constanza Aguado-Norese, Sylvain Prigent, Francisca P. Díaz, Dominique Rolin, Mauricio González, Caroline Müller, Rodrigo A Gutiérrez, Pierre Pétriacq

## Abstract

Plants can modulate their rhizosphere chemistry, thereby influencing microbe communities. Although our understanding of rhizosphere chemistry is growing, knowledge of its responses to abiotic constraints is limited, especially in realistic ecological contexts. Here, we combined predictive metabolomics with bacterial sequencing data to investigate whether rhizosphere chemistry responded to environmental constraints and shaped bacterial communities across an elevation gradient in the Atacama Desert. We found that metabolic adjustments of rhizosphere chemistry predicted the environment of four plant species independently of year, identifying important rhizosphere metabolic biomarkers. Inter-species predictions unveiled significant biochemical convergences. Subsequently, we linked metabolic predictors to variation in the abundance of operational taxonomic units (OTUs). Chemical response influenced distinct and common bacterial families between species and vegetation belts. The annotation of chemical markers and correlated bacterial families highlighted critical biological processes such as nitrogen starvation, metal pollution and plant development and defence. Overall, this study demonstrates a unique metabolic set likely involved in improving plant resilience to harsh edaphic conditions. Besides, the results emphasise the need to integrate ecology with plant metabolome and microbiome approaches to explore plant-soil interactions and better predict their responses to climate change and consequences for ecosystem dynamics.

## Introduction

The ability of a plant to thrive in an ecosystem is closely dependent on autonomous (*e.g.* genetic make up) and non-autonomous capacities (*e.g.* interactions with soil microbes) (1–3). Plants alter soil properties and drive soil microbiome and rhizosphere interactions via root exudates, a process called plant-soil feedback when it influences plant fitness (4–8). In turn, the soil microbiome influences the soil structure and can also positively or negatively influence plant growth (9–11). Besides, the interaction between plants and soil microorganisms was recently shown as a key driver of plant adaptation (12). The rhizosphere, which refers to the area around a plant root (13–15), is the nerve centre of intense chemical dialogue between the plant and the soil microbiome (6,14,16,17). Analysing rhizosphere chemistry is complex since it includes plant exudates, their breakdown products as well as microbial compounds (18). For instance, plants release organic acids to improve resource uptake or tolerance to soil toxicity (19,20). Plants exudate flavonoids, coumarins and lipids to condition rhizosphere microbiota and benzoxazinoids can be metabolised by microorganisms into antimicrobial compounds (7,8,14,21–23). Thus, analyses of non-sterile rhizosphere should be used as a holistic approach to capture a more complete fraction of rhizosphere signals, instead of plants isolated from their natural environment (18,24,25). Overall, recent efforts improved our understanding of some biochemical mechanisms used by plants to interact with the microbial community (26). Nevertheless, it is reasonable to say that a significant portion of rhizosphere chemicals remains to be discovered and that little is known about how these signals shape the surrounding microbe community (15,18,27). Moreover, most studies were performed on single species. Hence, rhizosphere chemicals should be explored in wild ecosystems and across multiple plant species to gain a more realistic image of soil chemistry *in natura*.

Soil microbe dynamics and plant-soil feedback are influenced by abiotic constraints (6,28,29). Previous studies detailed the effects of abiotic stress such as drought on mycorrhizal fungi, plant exudates and plant-microbe interactions (30–34). Interestingly, the response of soil chemistry to environmental pressures such as drought seems highly variable between plant species and environments (3). However, the plant species- or environment-specific character of most studies may explain this high specificity. Besides, most experiments were conducted under controlled conditions, which only enabled the analysis of short-term responses. In contrast, studies in wild ecosystems offer a unique opportunity to explore the responses of rhizosphere chemistry to abiotic stress that result from long-term adaptation processes.

The Atacama Desert is the driest non-polar desert on Earth (35). In this desert, the Talabre-Lejía transect (TLT) is an elevation gradient (≈2,500-4,500 m.a.s.l) characterised by extreme scarcity of water (20 to 160 mm.yr^-1^), nutrient deprivation, and high solar irradiance (600 W.m^-2^.d^-1^) (35). Tens of plant species thrive in this transect and define three plant communities: the Prepuna (2,500-3,300 m.a.s.l), the Puna (3,300-4,000 m.a.s.l) and the Steppe (4,000-4,500 m.a.s.l) (36). Thus, the Atacama Desert represents an ideal system to study responses of the rhizosphere chemistry to this gradient of extreme abiotic constraints. Besides, a high variation in nature and abundance of soil microorganisms has previously been reported for this area (29,35). Here, we collected four plant species and the associated non-sterile rhizosphere soil as well as associated bulk soil at various elevations to explore the responses of the rhizosphere chemistry to the different abiotic conditions. In addition, this biological diversity was used to investigate the specificity level of rhizosphere metabolic signals across plant species. Subsequent bioinformatic analyses linked rhizosphere metabolic predictors to the composition of the associated bacterial communities, based on operational taxonomic units (OTUs), across the gradient to gain an understanding of ecological dynamics.

## Materials and Methods

### Sampling

Non-sterile rhizosphere samples were collected in the Talabre-Lejía transect (lat 22°-24°S) in April (07–08) 2021 at 5 cm depth under four distinct plant species, whose aerial parts were also sampled (Tab. S1). Species included *Atriplex imbricata (Moq.) D.Dietr.* (Amaranthaceae), *Hoffmannseggia doellii* Phil. (Fabaceae), *Adesmia spinosissima* Meyen ex Vogel (Fabaceae) and the Poaceae *Jarava frigida* (Phil.) F. Rojas (synonym of *Pappostipa frigida*, (Phil.) Romasch). Rhizosphere samples under *J. frigida* and samples from *A. imbricata* and *J. frigida* were also collected in April (06–07) 2019 (Tab. S1). Each pair of samples (*i.e.* rhizosphere and associated plant) was collected in 1 to 4 elevation sites (Fig. 1), depending on species availability, with a minimum of three biological replicates. Two soil samples were taken from open areas (*i.e.* without plants above ground) within a ≈30 cm radius around the target species at each elevation. Samples were collected, directly snap-frozen into liquid nitrogen and transported to the laboratory in dry ice until freeze-drying and grinding. Dried sample powders were stored at -80°C until extraction.

**Fig. 1.**
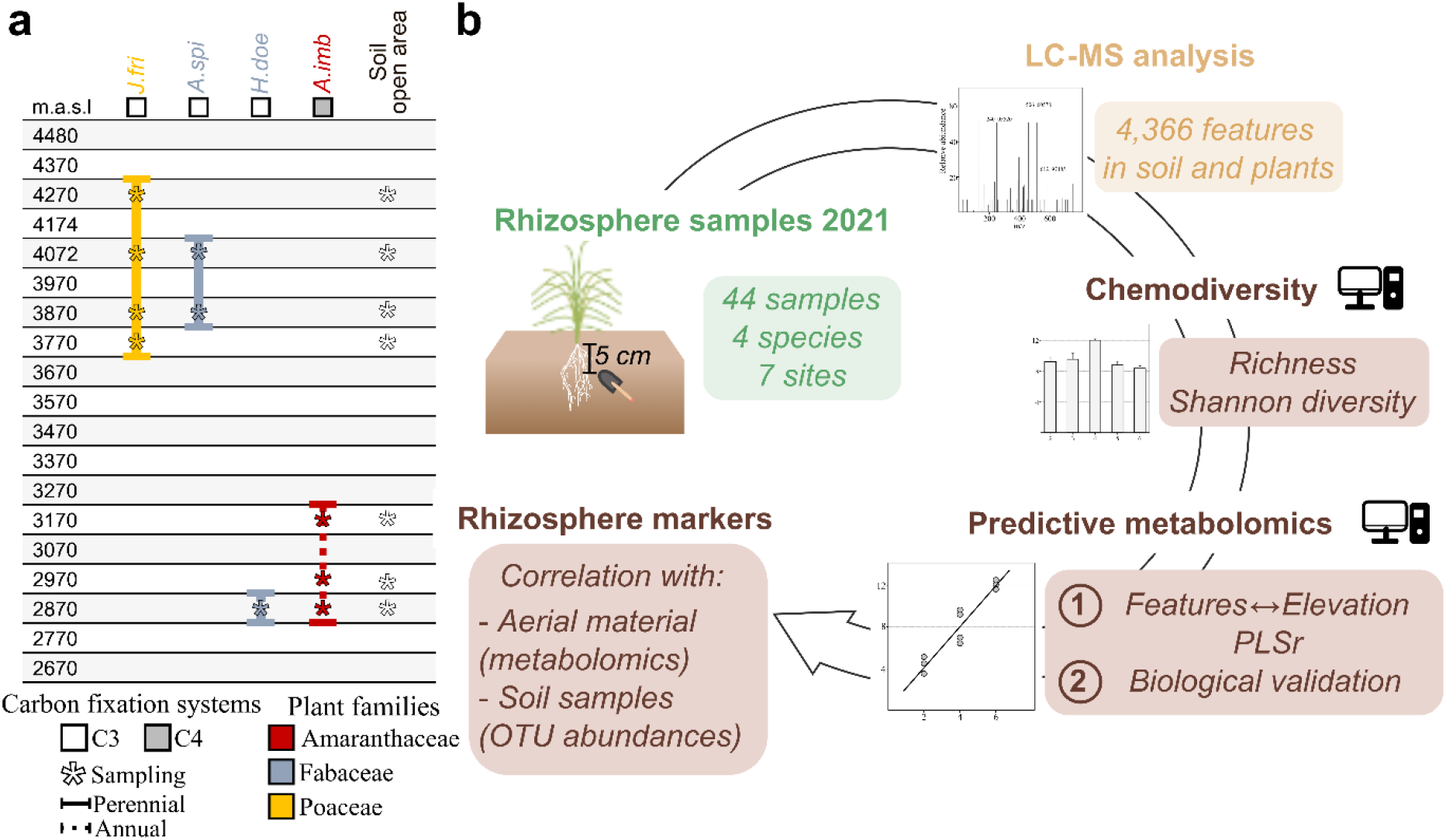
Sampling and analytical workflow. **a.** Sampling of plant and rhizosphere material. **b.** Simplified workflow of the analysis. *A.imb*: *Atriplex imbricata*, *A.spi*: *Adesmia spinosissima*, *H.doe*: *Hoffmannseggia doellii*, *J.fri*: *Jarava frigida*.

### Environmental data

Climatic data were assessed using two meteorological stations (3,060 and 4,090 m.a.s.l) and soil composition was previously obtained as described (35).

### Metabolomics

To maximise the extraction of semi-polar compounds, we performed robotised ethanolic fractionation using 450 mg or 20 mg of dried soil and plant material, respectively (37). Ethanol extracts were subjected to ultra-high-pressure liquid chromatography (UHPLC) - Orbitrap mass spectrometry analysis using an Ultimate 3000 UHPLC combined with an LTQ-Orbitrap Elite MS (ThermoScientific, Bremen, Germany). The separation was realised on a C18 column (C18-Gemini, 150 x 2 mm, 3 µm, 110 Å, Phenomenex, France) coupled to a C18 SecurityGuard Gemini pre-column (4 × 2 mm, 3 µm, Phenomenex, France) at 30°C with an injection volume of 5 µL, a flow rate of 350 µL/min and a previously established gradient (37). The LTQ-Orbitrap was operated in negative electrospray ionisation (ESI^-^) mode (37,38). MS spectra were acquired using data-dependent analysis at a resolution of 30,000 in a *m/z* range of 50-1,500. Quality controls (QC) were injected every 10 samples.

Raw LC-MS data were processed via MS-DIAL (v. 4.90) using optimised parameters (39) detailed in Table S2. Putative annotation was performed using the MS/MS-Public-Neg database (v. 17). Feature filtering was performed using blanks and QC samples, where features exceeding a coefficient of variation of 30% were excluded. The resulting dataset (4,629 features) was first normalised by sample weight, followed by median normalisation, cube-root transformation and Pareto scaling using MetaboAnalyst (v. 3) (40) before statistical analyses. The non-normalised dataset obtained after preprocessing is available in Table S3 and was deposited online (see Data availability).

### Statistical analyses

The normalised dataset was processed through multivariate analysis on MetaboAnalyst (v. 3.0) and R software (v. 4.2.1) (40,41). Principal component analysis (PCA) was performed using *FactoMineR* package (42). Variation in chemical diversity was explored using the *chemodiv* package to calculate richness and Shannon’s indices (43). To extract rhizosphere or plant features responding significantly to elevation, we performed ANOVA tests on MetaboAnalyst and tested Pearson’s correlation using *Hmisc* package (44). Features responding significantly to elevation and significantly correlated with elevation (Pearson’s correlation, *P* < 0.05, FDR correction) were extracted for subsequent analyses. To test the capacity of rhizosphere chemistry to predict plant elevation, we performed partial least squares regression analyses (PLSr) via the *pls* package, which was used to select the optimal number of components (“onesigma” function, one component was selected) and to perform the predictions (45). Sample sets were divided into a “training” (80%) and a “testing” set (20%) using stratified sampling as previously described (38). Models were performed 50 times to cope with random partitioning. To test the likelihood of spurious predictions, 50 permutation sets were created for each model by randomly swapping elevation levels between samples. Next, the predictive capacity of the predictive features (hereafter referred to as markers) was confirmed by performing real predictions on independent datasets. Model equations were developed on plants (*A. imbricata* or *J. frigida*) collected in 2021 and directly applied to either (i) the other two species (*H. doellii*, *A. spinosissima*) and (ii) plants of the same species collected in 2019. Tukey’s tests were performed using the *agricolae* package (46). Figures were designed using *ggplot2*, *ggpubr* and *Hmisc* packages (44,47,48).

### Annotation and classification

The automatic annotation (see above) was complemented by a manual putative annotation for the best predictive features as previously described (38). Putative chemical formulas were defined by screening accurate *m/z* on MetFrag (49), and the putative annotation was performed using ChEBI (50) and KNApSAcK (http://www.knapsackfamily.com/KNApSAcK/) databases. MS/MS spectra were compared to experimental spectra on Massbank (51). The metabolomics standards initiative levels (MSI levels) were used to define the confidence level in the putative annotations (52), as previously performed (38,53). All annotations should be considered putative, even when confirmed with an MS/MS match (MSI 2 level). Biochemical classes and pathways were defined using ChEBI and ClassyFire taxonomy (54).

### Correlation of the rhizosphere markers with aboveground metabolism

To link rhizosphere metabolic predictors with aboveground metabolism, we first extracted metabolic features that responded significantly to elevation in aboveground samples of *A. imbricata*, *A. spinosissima* and *J. frigida* collected in 2021 and 2019 (Tukey’s or Student’s test, *P* < 0.05, FDR correction). We then tested whether rhizosphere metabolic predictors were also responding significantly to elevation in aerial tissues (Fig. S1).

### Correlation of the rhizosphere markers with soil microbiome

Next, the intensity of the rhizosphere markers was linked to changes in the composition of the soil bacterial community (Fig. S1). Previously, the relative abundance of OTUs had been assessed in both bulk soil (BS) and rhizosphere surrounding samples (RSS) under multiple plant species, each at a given altitude, through the Atacama transect (29,35). For each dataset (BS and RSS), OTU data were rarefied to 8000 reads and only OTUs identified at the family level were used for our analysis (10,515 OTUs). We extracted OTUs responding significantly to elevation (*P* < 0.05, FDR) and detected in at least three RSS samples to exclude OTUs found under specific plant species or genera, yielding a set of 1,973 OTUs. To test for convergent responses of microbial communities to elevation, PLSr models were performed as detailed above (see Statistical analyses). Real predictions were performed by excluding four species from the RSS dataset (*Exodeconus integrifolius*, 2,870 m.a.s.l; *Tricholcline caulescens*, 3,370 m.a.s.l; *Lupinus subinflatus*, 3,870 m.a.s.l; *Calamagrostis cabrerae*, 4,270 m.a.s.l). Model equations were developed on the remaining dataset and directly applied to these four species to predict their elevation. The most predictive OTUs (top 1%, 105 OTUs) were defined based on their coefficients in the models. Putative functional roles of the top 1% predictive OTUs were assigned via the *PICRUSt2* package (55) using 16S rRNA sequences and the KEGG database as previously described (29).

Next, we linked the response of plant rhizosphere chemistry and bacterial communities. According to our results, convergent responses of plant rhizosphere to elevation might be observed, which could lead to similarities in the response of certain microbial species. In this context, correlations between OTUs abundances and metabolic intensities were tested regardless of the plant species under which samples were collected. For Steppe species, the average intensity of the best 216 markers was calculated for each elevation at which *J. frigida* or *A. spinosissima* were collected in both experiments (*i.e.* 3,870, 4,072, 4,270 m.a.s.l, Fig. S1).

OTUs abundances of RSS samples at the same elevations were extracted (30 samples, 10 plant species, 3 elevations) and Pearson’s correlation was assessed. We defined a significant correlation between a metabolic marker and an OTU if *P* < 0.05 after FDR correction. Similarly, the average intensity of the best markers from Prepuna species (*A. imbricata* and *H. doellii*) at 2,870 and 3,170 m.a.s.l was linked to OTU abundances from RSS samples collected at similar elevations (9 samples, 3 plant species, 2 elevations, Fig. S1). The most relevant bacterial families were defined by exploring the total number of related OTUs correlated to at least one metabolic marker. A literature search was carried out to develop hypotheses about the potential roles of these microbes.

## Results

### Influence of plant species and environment on rhizosphere chemistry

To study how rhizosphere signals respond to an extreme stress gradient, we collected aerial parts of four plant species and their associated rhizospheres at different elevations from 2,870 to 4,270 m.a.s.l. in the Atacama Desert (Fig. 1a and Tab. S1). Untargeted eco-metabolomics was deployed to assess rhizochemicals (Fig. 1b). LC-MS analyses yielded 9,008 features after pre-processing, from which 4,629 were detected in rhizosphere and 4,366 in both rhizosphere and aboveground samples (Tab. S4). As a first step, the focus was placed on the metabolic features detected in both rhizosphere and plant samples. In an unsupervised analysis of metabolic fingerprints of the non-sterile rhizospheres, plant communities (*i.e.* Prepuna and Steppe species) were split across the second component while plant species and elevation factors mediated chemical variation within a given community along the first or second component (for a total variance of 35.7%) (Fig. S2). Hence, this PCA suggested an influence of both (i) plant species and (ii) environment on rhizosphere chemicals.

### Chemical diversity decreases under challenging conditions

Next, we investigate the impact of plant species and environment on rhizosphere signals. Richness and Shannon’s diversity of the rhizosphere chemistry were highly dependent on plant species and elevation (Fig. S3). Prepuna species (*A. imbricata* and *H. doellii*) displayed lower metabolite numbers when compared to Steppe species (*A. spinosissima* and *J. frigida*). Chemical richness decreased in highly stressful environments (*i.e.* lowest and highest elevations). An impact of elevation was also observed when defining chemodiversity indices for rhizosphere samples collected at 3,770 m.a.s.l under *J. frigida*, which were comparable to that observed in lowland species (Fig. S4c and S4d). Within species, chemical richness decreased at the edges of the life-compatible gradient (Fig. S4). For instance, the lowest richness scores were detected at 3,770 and 4,480 m.a.s.l in the rhizosphere of *J. frigida*. Unfortunately, *A. imbricata* was not sampled at its lowest and highest survival sites, explaining the distinct pattern for this species (36). Shannon indices of the rhizosphere chemical pattern were likewise lower at the edges of the elevation gradient, but to a lesser extent (Fig. S3 and S4). These results suggest that plants or microbes tend to narrow their chemical repertoire under stressful conditions.

### Rhizosphere chemistry predicts plant environment

We then tested the response of rhizosphere chemicals to the extreme elevation gradient in *J. frigida* and *A. imbricata*, which were collected on a consequent elevation delta (500 and 300 m, respectively). In *J. frigida*, 216 features detected in rhizosphere and plants were linked to elevation (Tukey’s test and Pearson’s correlation, *P* < 0.05, FDR correction). Similarly, 8 features responded significantly to elevation in *A. imbricata*, while 54 showed a significant trend (*i.e. P* < 0.05 without FDR correction). In comparison, 5 significant features detected in rhizosphere and aerial parts of *A. spinosissima* (Student’s test, *P* < 0.05, FDR correction), which was collected at only two elevations with a 200 m delta (Tab. S5), were observed. To link plant environment and rhizosphere chemistry, we investigated whether rhizosphere chemicals could predict plant elevation, which was used as a proxy of the environment. Sample sets from *A. imbricata* and *J. frigida* collected in April 2021 were divided into a training (80%) and a testing set (20%) as previously described (38). For each species, 50 PLSr models were developed and prediction quality was assessed by fitting predicted elevations to measured values. In *A. imbricata*, rhizosphere chemistry predicted elevation level with 0.96 and 0.91% accuracy using 8 or 54 features, respectively (Fig. 2 and S5). Rhizosphere chemistry predicted plant environment with an average *R²* of 0.74% in *J. frigida*. A ranking of the metabolic features was proposed based on their coefficient in the models (Tab. S6). Besides, the randomness of these predictions was excluded using 50 permuted sets for each species (Fig. 2). The predictive capacity of rhizosphere chemistry was also observed in *A. spinosissima*, where the 5 significant features discriminated elevation levels with 0.99% accuracy (Fig. S5). Subsequently, the robustness of the predictive features (hereafter referred to as markers) was tested on *J. frigida* by using an independent dataset composed of rhizosphere samples under *J. frigida* collected in April 2019. The model equation was defined on samples collected in April 2021 and directly applied to rhizosphere samples collected in April 2019 to calculate their elevation levels, resulting in a similar predictive capacity (*R²* = 0.79%) (Fig. 2). Importantly, the linear response of rhizosphere chemicals under *J. frigida* was supported by the accurate prediction of rhizosphere samples collected at 4,480 m.a.s.l, an elevation that was not present in the 2019 sample set. Model predictions were validated using 50 permutation sets (Fig. S5).

**Fig. 2.**
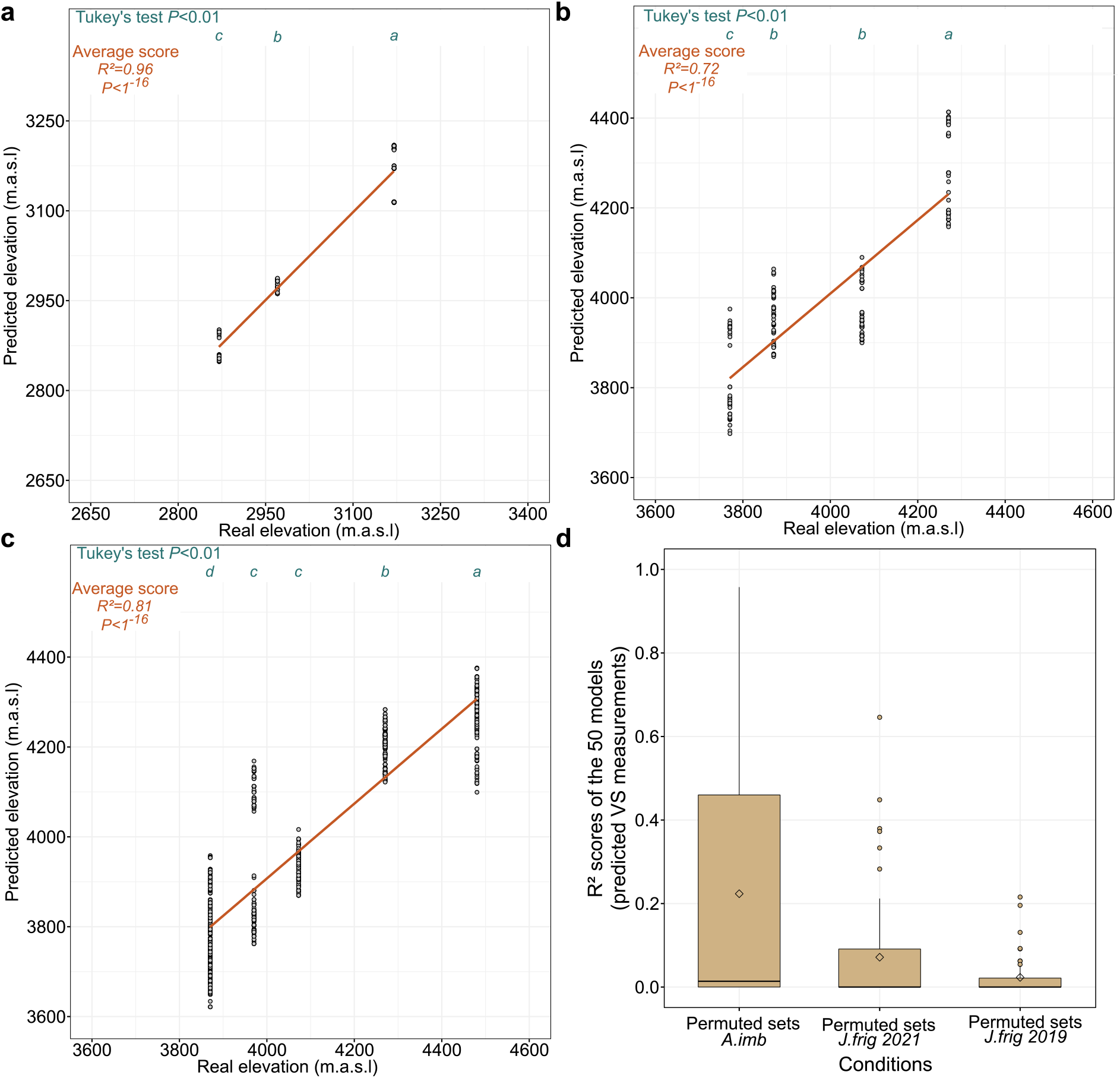
Prediction of elevation levels using metabolic fingerprints of rhizosphere samples. **a.** Partial least squares regression (PLSr) analyses using *A. imbricata* significant features (8 features). **b.** PLSr analyses using significant features (216 features) of rhizosphere samples collected under *J. frigida* in 2021. **c.** PLSr analyses of rhizosphere samples collected under *J. frigida* in 2019. Model equation was developed on samples collected in 2021 and directly applied to 2019 rhizosphere samples (216 features). **d.** PLSr analyses using 50 permuted datasets. In boxplots, the diamond represents the mean while the solid dark line represents the median. PLSr models were performed 50 times. Tukey’s tests were performed to compare predictions between elevation levels (*P* < 0.01).

To test whether soil-specific features (*i.e.* features detected in soil samples exclusively) could improve predictive scores, we conducted the same analytical workflow. Although some soil-specific features responded significantly to elevation (*P* < 0.05, FDR correction), predictions were not improved when adding these features to the PLSr models (Tab. S7 and S8). Altogether, these results demonstrate that variations in environmental conditions induce a significant change in rhizosphere chemistry, which could be used to deduce the altitude of the sample independently of the year of sampling.

### Potential convergent response of rhizosphere chemistry

Putative annotation of the significant metabolic markers revealed that in the rhizosphere of *A. imbricata*, secondary (or specialised) metabolism was predominant (63%) (Fig. 3a and Tab. S9). Phenolics such as flavonoids were the most widely represented chemical metabolite class, confirming their essential role in the adaptation of Atacama plants (38). Furthermore, terpenes, coumarins and cinnamic acids were included among metabolic predictors. Primary compounds such as organic acids and compounds derived from purine and pyrimidine pathways were also reported (10 and 3 compounds, respectively). For *J. frigida* rhizosphere, putative annotations yielded a certain equilibrium between primary (42%) and secondary (57%) metabolism, which was mainly explained by the overrepresentation of lipids (Fig. 3b). While the significance of metabolites from the phenolic pathways was again confirmed, terpenes were the most represented metabolites in this species. Triterpenoids, sesquiterpenoids but also derivatives such as terpene glycosides were detected among the markers. Within primary metabolism, a high proportion of the predictors referred to lipids (40 compounds), while likewise organic acids were present. Overall convergent and divergent responses were observed depending on the plant species under which rhizosphere samples were collected. To test the convergence degree, we tried to predict the rhizosphere elevation level under different species using samples of *A. imbricata* or *J. frigida*. The elevation of rhizosphere samples from *H. doellii* and *A. spinosissima* was predicted using the equation developed on *A. imbricata* or *J. frigida*, respectively. The elevation of both species was considerably predictable with accurate predictions of *H. doellii* elevation and an *R²* of 0.74 for predictions of *A. spinosissima* (Fig. 4). This result suggests convergent chemical responses of the surrounding rhizosphere to the elevation gradient between species.

**Fig. 3.**
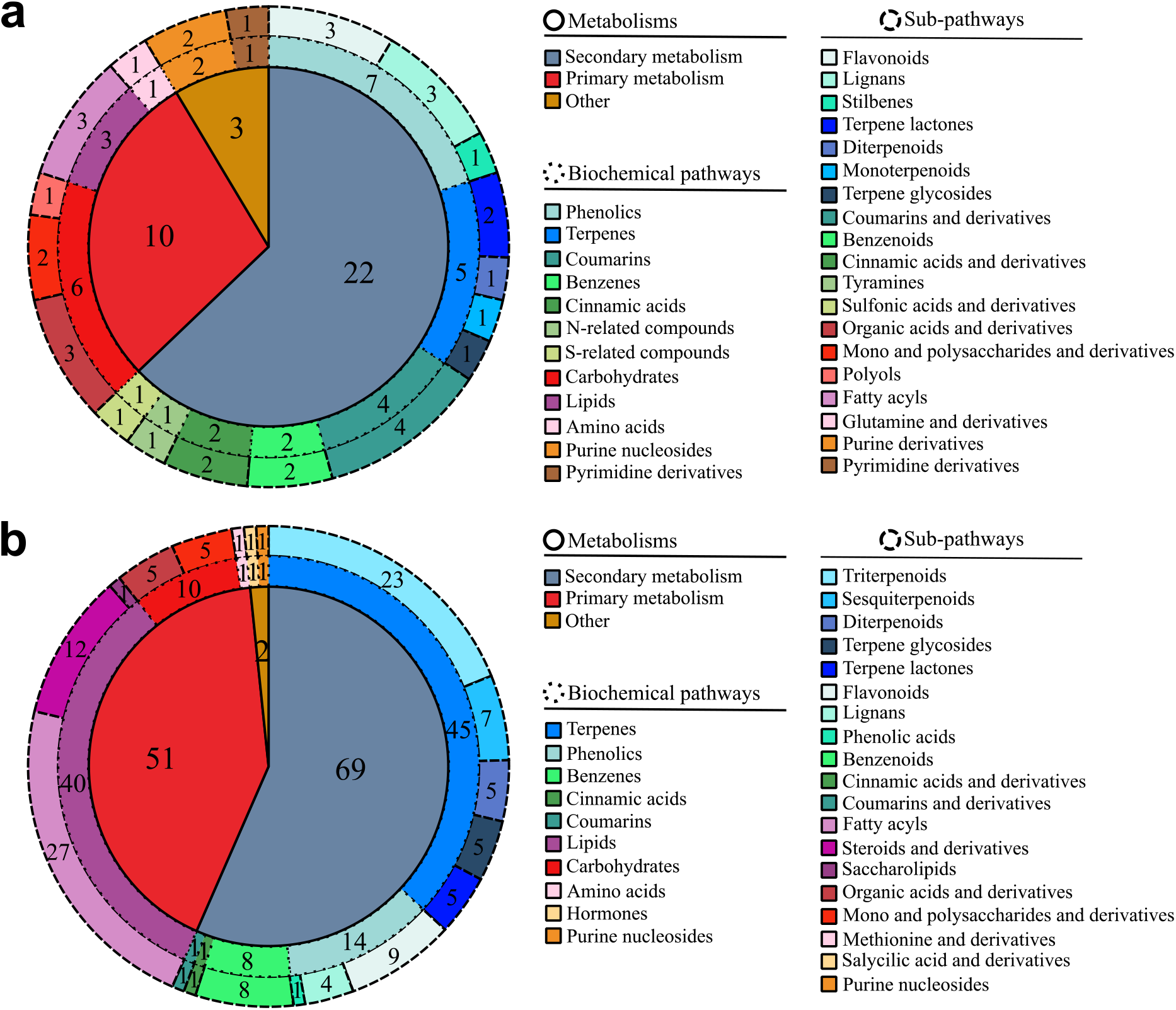
Annotation of the metabolic markers. **a.** Annotation of elevation markers from *A. imbricata*. **b.** Annotation of elevation markers from *J. frigida*. Unknown metabolic features were not displayed in this figure. See Table S9 for more details.

**Fig. 4.**
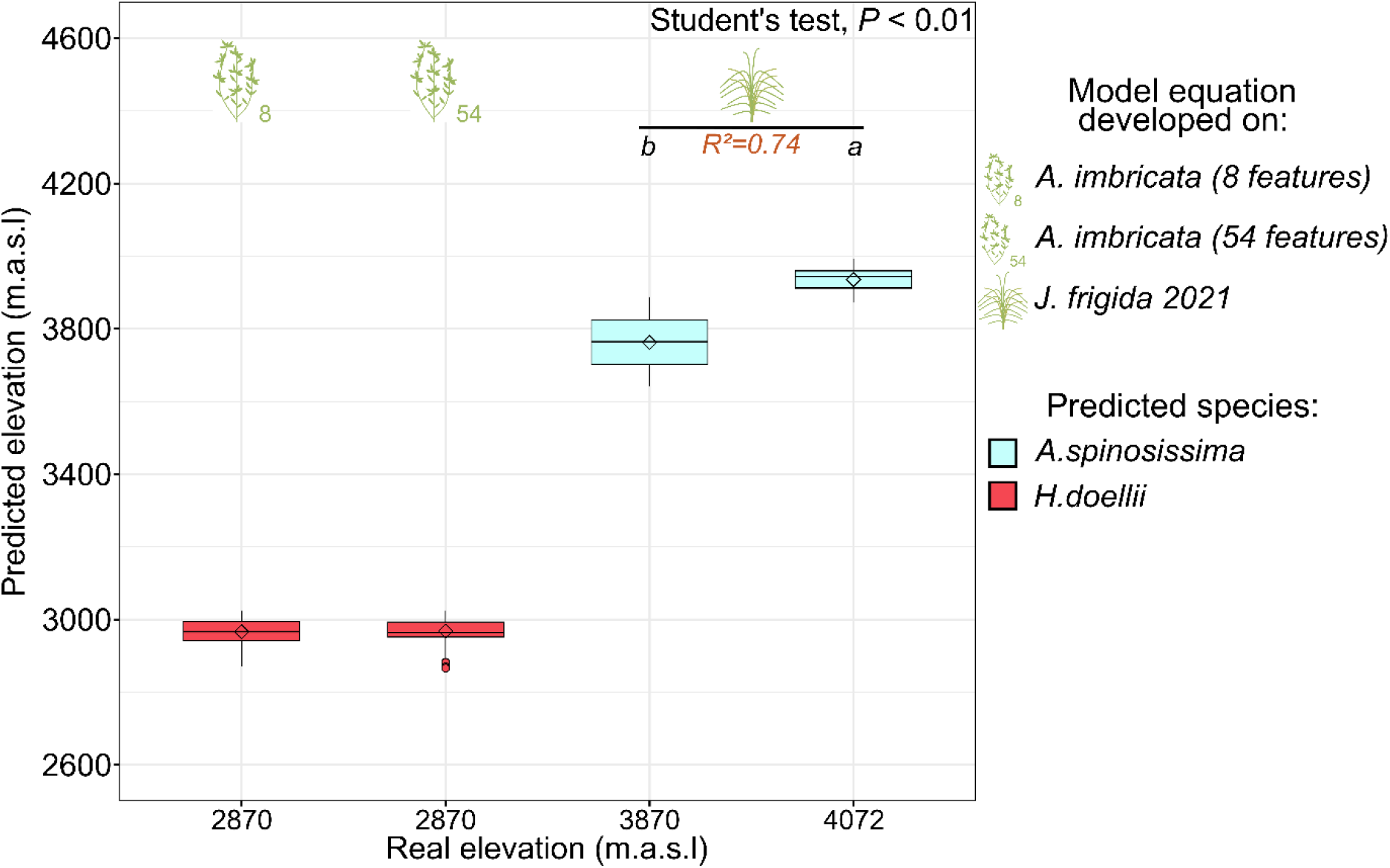
Prediction of elevation levels from independent conditions using metabolic markers of *A. imbricata* or *J. frigida*. Partial least squares regression analyses (PLSr) to predict elevation levels of *H. doellii* and *A. spinosissima*, respectively. Model equations were developed on *A. imbricata* (8 or 54 features) or *J. frigida* and directly applied to the independent samples. PLSr models were performed 50 times. Student’s test was performed to compare predictions between elevation levels (*P* < 0.01). In boxplots, the diamond represents the mean while the solid dark line represents the median.

### Convergences are also found in the response of microbial communities to the elevation gradient

To test whether convergences could reflect or be also observed in the response of soil bacterial communities, we used previous data on variation in OTU abundances across the transect for 21 plant species (29,35) (Fig S1). While these analyses deciphered the variation in bacterial families between species, we asked whether OTU abundances could be used to predict elevation levels independently of the species under which rhizosphere samples were collected. In the initial OTU datasets (10,515 OTUs) of rhizosphere collected from 21 plant species, we extracted OTUs detected in at least 3 samples to exclude OTUs found under specific plant species or genera. From this set, 1,973 OTUs responded significantly to elevation (*P* < 0.05, FDR). PLSr models were first deployed on the entire set to verify the capacity of OTU abundances to predict elevation levels, yielding an average *R²* of 0.80 for 50 fits (Fig. S6). “Real” predictions of elevation were performed for 4 species (2,870, 3,370, 3,870 and 4,270 m.a.s.l) previously excluded from the dataset, yielding an average *R²* of 95%. The top 1% predictive OTUs (*i.e.* 105) allowed for similar predictive scores and models were validated using permuted datasets (Fig. 5a, Tab. S10). From these 105 OTUs, the most represented taxon positively correlated to elevation levels was Gaiellaceae. In contrast, Rubrobacteraceae and Cytophagaceae were the most negatively correlated taxa. In addition, some microbe families such as Bradyrhizobiaceae, Sphingomonadaceae and Hyphomicrobiaceae were found either positively or negatively correlated to elevation, suggesting various roles depending on environmental demands (Fig. 5b). A functional analysis was then performed on the most predictive OTUs to depict the underlying putative KEGG Orthology (ko) pathways (Tab. S11). Great overlaps were observed between the annotation of the most predictive OTUs and the predictive rhizosphere chemicals. For instance, pathways linked to nitrogen metabolism, synthesis of precursors (*e.g.* amino acids) and secondary metabolism (*e.g.* terpenoids) were observed in both analyses, while others were overrepresented (*e.g.* pentose phosphate pathway, glycolysis) or exclusive to OTU analysis (*e.g.* glutathione and ascorbate metabolism) (Tab. S12). Next, we explored the place of the 1% most important predictive OTUs in the network of microbial interactions developed previously for the whole transect (29). Predictive OTUs represented a great proportion of the most correlated OTUs (*i.e.* highest degree centrality and drivers defined by NetShift analysis), highlighting their significance in the dynamics of microbial communities. Besides, the positive ratios of positive:negative connections (*i.e.* 1.7 and 3.8 for Steppe and Prepuna, respectively) suggested a need for cooperation at the most extreme elevations, which correspond to the most stressful environments (Tab. S13). Altogether, these results shed light on the existence and significance of chemical and ecological (*i.e.* dynamics of microbe communities) convergences belowground in response to the elevation gradient.

**Fig. 5.**
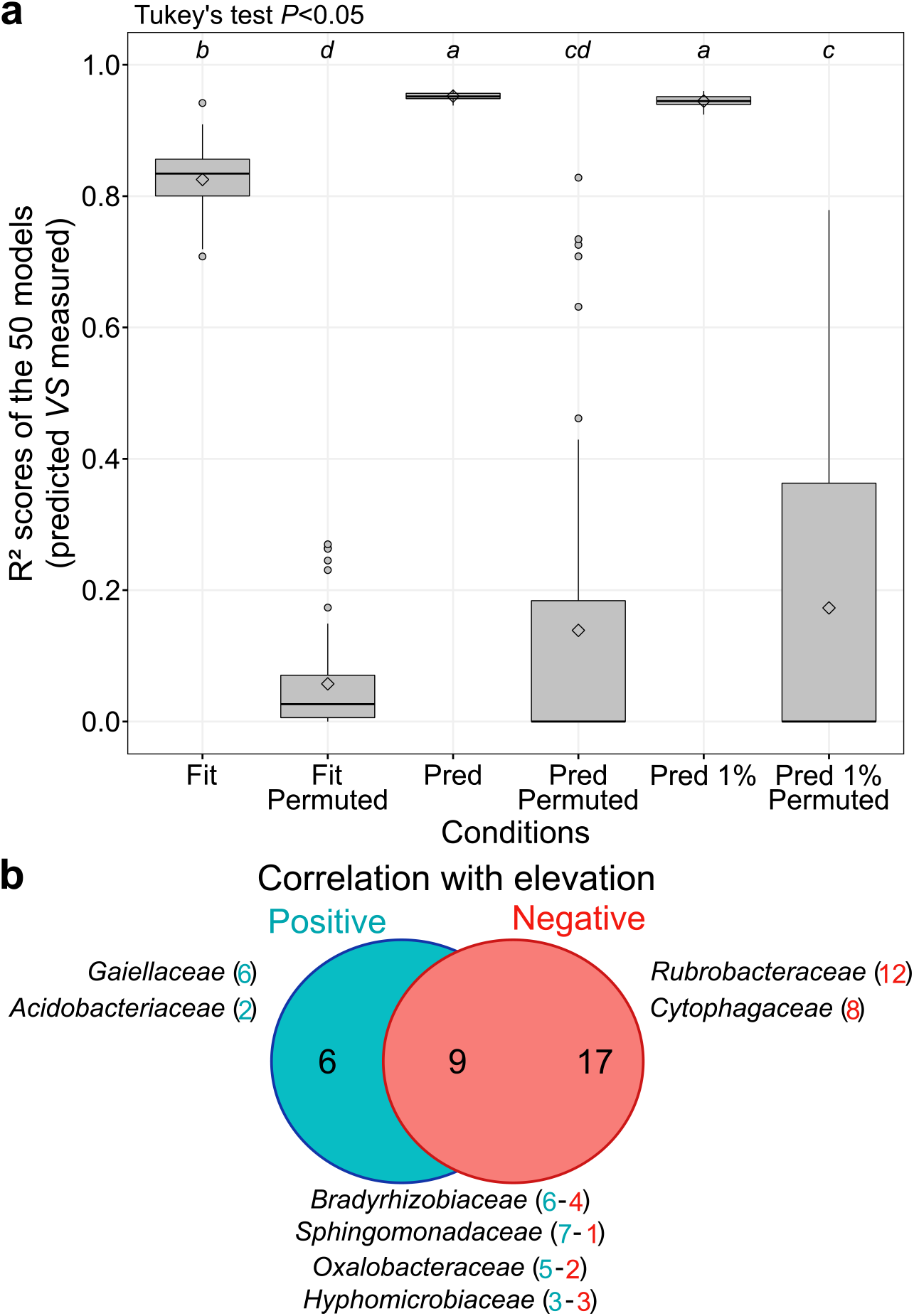
Response of soil bacterial communities to the elevation gradient. **a.** Predictive capacity of PLSr models using significant OTUs (*Fit/Pred* models, 1,973 OTUs) or the top 1% most predictive OTUs (*Pred 1%* models, 105 OTUs) to predict the elevation of rhizosphere samples. All samples were used for “Fit” models. For predictive models (*Pred*), four species were excluded from the dataset (*i.e.* testing set). Model equation was developed on the remaining samples and directly applied to the testing set. Fifty permuted datasets were developed for each modelling condition. Tukey’s tests were performed to compare predictions between elevation levels (*P* < 0.05). **b.** Venn diagram of the bacterial families involved in the top 1% predictive OTUs. Numbers in parentheses refer to the number of OTUs included in the corresponding bacterial family. While examples are provided, more details are provided in Tab. S10.

### Rhizosphere chemical markers are linked to the response of aboveground metabolism and microbe communities

We tested the relationship between the rhizosphere predictive metabolites (*i.e.* markers) predicting plant environment and the response of both aboveground metabolism and soil bacterial communities to elevation (Fig. S1). A total of 19 (32% of *A. imbricata* rhizosphere markers) and 64 (30% of *J. frigida* rhizosphere markers) markers were also significant in aboveground samples from at least one plant species (*P* < 0.05, FDR, Fig. 6a, Tab. S14). Without FDR correction, these numbers reached 63% and 58% for *A. imbricata* and *J. frigida* markers, respectively (Tab. S15). Some markers were significant in leaves from two or three species. When linking the best metabolic predictors to microbial communities, potential convergent responses of rhizosphere chemistry and bacterial communities were noticed. Hence, correlations between OTU abundances and the intensity of the best metabolic markers were tested independently of the species under which rhizosphere samples were collected (Fig S1). Very high correlations were observed (Fig. 6). A total of 111 OTUs (10 species, 3 elevations) were correlated with the average intensity of at least one *J. frigida* rhizosphere marker, including 34 microbe families (Pearson’s correlation, *P* < 0.05, FDR). Similarly, 184 OTUs (3 species, 2 elevations) were correlated with *A. imbricata* rhizosphere markers, which included 42 bacterial families (Table S16 and S17). Interestingly, great overlaps were observed with 21 families that included at least one related OTU correlated to a metabolic marker (Fig S7). In addition, these bacterial families strongly referred to the top 1% predictive OTUs used to predict elevation levels. A literature survey showed that overrepresented bacterial families were associated with potential effects on litter decomposition, nitrogen cycle and metal detoxification (Fig. 6b and 6c and Tab S17 for references). Variation in nitrogen fixers across the elevation gradient (*e.g.* Bradyrhizobiaceaee, Hyphomicrobiaceae, Acetobacteraceae) was linked to chemical markers in both Prepuna and Steppe species. Gaiellaceae, which are known for their implications in the phosphorous cycle, were also observed in both vegetation belts. In contrast, growth-promoting bacteria (*e.g.* Geodermatophilaceae, Sphingomonadaceae) and bacteria linked to metal detoxification (*e.g.* Caulobacteraceae) were more represented in Prepuna. Micrococcaceae were exclusive to Prepuna, where salts are accumulated.

**Fig. 6.**
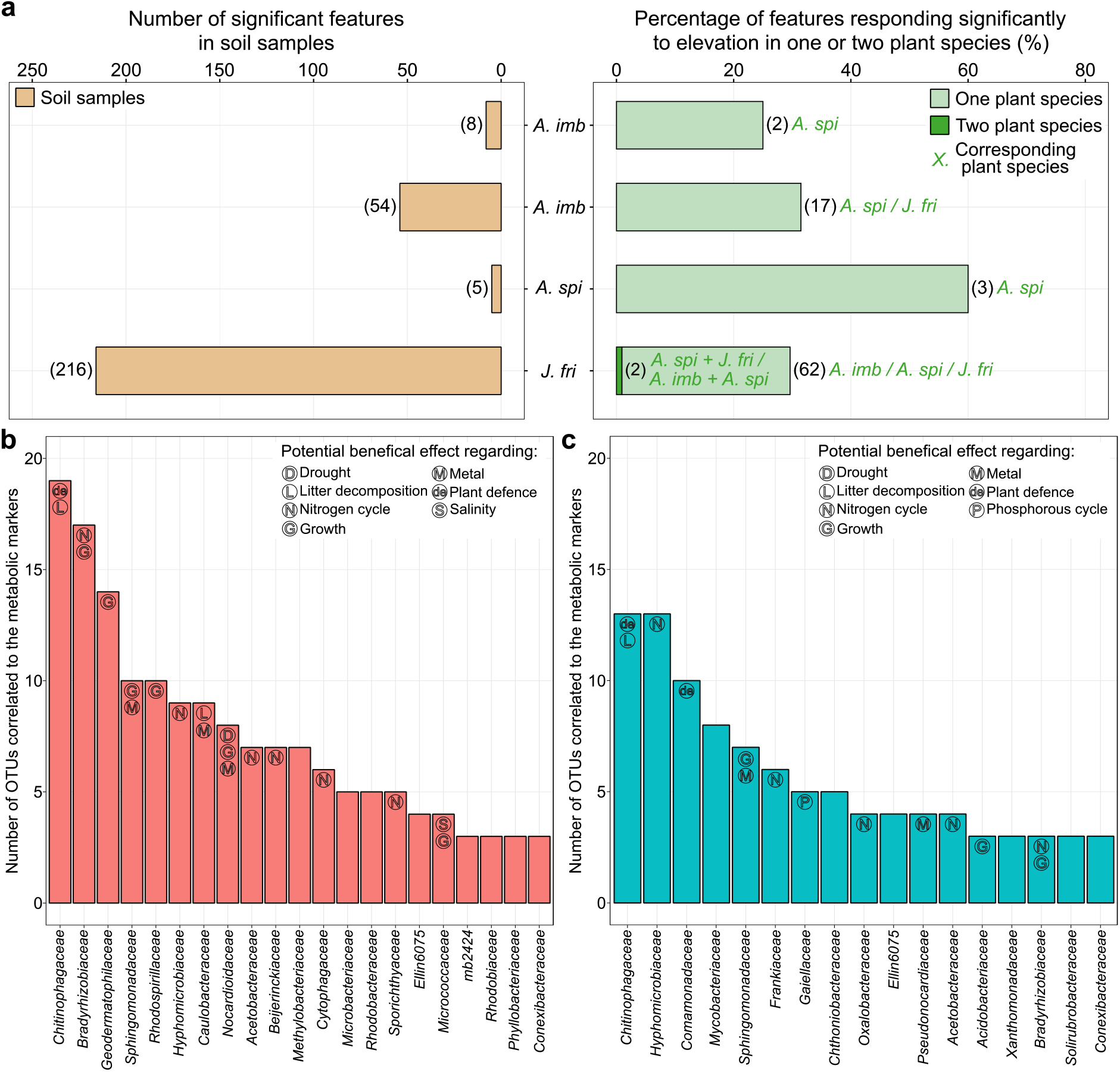
Correlations between rhizosphere metabolic predictors and plant aboveground response and soil microbiome. **a.** Numbers in brackets indicate the number of significant features in rhizosphere or plant samples for a given condition. All reported plant features were significant using Tukey’s test (P<0.05, FDR correction). *A. imb*: *A. imbricata*, *A. spi*: *A. spinosissima*, *J.fri*: *J. frigida*. **b-c**. Correlated microbe families to the best metabolic markers (*P* < 0.05, FDR correction, Pearson’s correlation) from *A. imbricata* in the Prepuna (**b**) or *J. frigida* in the Steppe (**c**). For ease of view, bacterial families were presented only if at least three related OTUs were significantly correlated. Additional information in Tab. S16 and S17.

Finally, we tested whether a correlation between chemical markers and OTU abundances could be observed in bulk soil samples (BS). In this context, we extracted the OTU abundances from BS following the same protocol as for RSS (Fig S1). A total of 647 and 14 BS OTUs were correlated (*P* < 0.05, FDR) with at least one metabolic marker from *J. frigida* and *A. imbricata* rhizosphere samples, respectively (Tab. S18). To test whether diffusion of the rhizosphere chemical response could explain such differences, we tried to predict the elevation level of soil samples in open areas using the equation of either *A. imbricata* or *J. frigida*. Interestingly, while *J. frigida* rhizosphere could be used to predict the elevation of samples collected in open areas with 64% accuracy, a similar trend was not observed (*R²=*0%) when using *A. imbricata* rhizosphere (Fig. S8). Overall, by combining predictive metabolome and soil microbiome analyses we could highlight that (i) rhizosphere chemistry captures environmental variation in extreme ecosystems, (ii) convergent and divergent chemical responses are observed between plant species and environments, and (iii) that these responses are potentially associated with the dynamics of soil bacterial communities.

## Discussion

### Rhizosphere chemistry is tailored to natural extreme environments

The importance of plant-soil feedback as a driver of ecosystem dynamics has been largely recognised (56,57). Studies have shown that rhizosphere chemistry is impacted by abiotic perturbations (58), but most of these studies were performed in controlled environments, lacking the ecological context and relevance of such variation. Here, we applied predictive metabolomics on rhizosphere samples under four plant species in the extreme Atacama Desert to uncover that rhizosphere chemistry is tailored to environmental constraints. First, significant variation was observed in chemodiversity indices. Richness and Shannon’s diversity were lower at low elevations, which coincided with the lifespan variation, with mostly annual plants inhabiting the Prepuna (2,400-3,300 m.a.s.l) and perennials the Steppe (4,000-4,500 m.a.s.l). These results are consistent with the effect of abiotic constraints on chemodiversity indices and illustrate the stress gradient faced by Atacama species (35,59). In addition, each of the four investigated plant species thrives on its specific elevation gradient and negatively modulates the richness of its rhizosphere chemistry when environmental pressure becomes critical, *i.e.* at the upper or lower altitude limits at which the species survives. However, while reducing their chemical repertoire, plants must maintain the regulation of specific compounds to thrive in these extreme altitudes, which should allow for predictions of elevations based on rhizosphere chemistry. This hypothesis was supported by the ability of several rhizosphere metabolites to predict the elevations of *A. imbricata* and *J. frigida* with 96 or 74% accuracy, respectively.

### Complementary metabolic strategies in below- and aboveground tissues to face harsh climatic and edaphic constraints

Most rhizosphere metabolic markers predicting plant environment in the present study were linked to nitrogen and phosphorous cycles, salt excess or water deprivation, as well as soil microbiome. For instance, markers included allantoin, guanosine and glutamic acid, which relate to the nitrogen cycle and purine degradation pathways (60,61). Organic acids (*e.g.* succinic acid), including ursolic acid, were previously linked to soil quality and salt stress mitigation (19,62). Terpenes, flavonoids, coumarins and cinnamic acids were highlighted in both vegetation belts, in agreement with their functions at the plant-microbe interface (8,63). Additional markers, such as lipids, were mainly observed in the rhizosphere of *J. frigida*. Lipids are considered as key drivers of plant-microbe interactions (64). A significant portion of these rhizosphere markers also responded significantly to elevation in aboveground plant material. These observations suggest that many of the rhizosphere markers were likely the result of plant responses to the environment rather than microorganism-derived processes. Overall, these results are complementary to previous studies. For example, transcriptomic analysis of 32 Atacama species identified positively selected genes closely linked to resource acquisition (35). In contrast to our rhizosphere analysis, a metabolomics study on leaves of Atacama plants uncovered compounds protective against extreme temperature, solar irradiance and drought, but very few metabolites linked to nutrient deprivation (38). Hence, the survival of Atacama plants may lie in their ability to adapt an organ-specific metabolic strategy to cope with severe climatic and edaphic constraints, from leaf to root, in coordination with their microbiome.

### Convergence in metabolic and ecological adaptation to the extreme environmental gradient

Studies have shown a strong phylogenetic signal in plant-soil feedback, while metabolic convergence was observed in aerial plant parts in response to abiotic threats (35,38,65,66). However, root and root exudate metabolic profiles tend to present less variation between species when compared to aerial plant parts (15). Here, while certain markers differed between vegetation belts (*e.g.* lipids), our results suggest a certain convergence in adaptations to the environment in Steppe or Prepuna species. In addition, the rhizosphere markers predicting the environment of *J. frigida* or *A. imbricata* were able to substantially predict the elevation of *A. spinosissima* (74% accuracy) and *H. doellii*, respectively. The analysis of OTU abundances also provided insights into the existence of convergent patterns in the dynamics of microbe communities across the elevation gradient. In line with the fact that extreme lands are recognised as hotspots for adaptive convergence (67), our results suggest that these events may not be restricted to aerial tissues. While this discovery represents a major advance in the field, further analyses comparing the response of rhizosphere chemistry under a wider range of species would be needed to support this hypothesis.

### Chemical rhizosphere markers are linked to growth-promoting bacteria

Associations between plants and microorganisms are vital to allow plants to survive in extreme ecosystems (29,57,68). Rhizosphere markers predicting plant elevation were significantly correlated with OTU abundances, highlighting the influence of rhizosphere chemicals on soil bacterial communities (14,32,69). OTU of the Chitinophagaceae were the most correlated microbes in both Prepuna and Steppe environments, reflecting the need to facilitate litter decomposition and defence against soil pathogens (70,71). Hyphomicrobiaceae, Bradyrhizobiaceae and Acetobacteraceae, which are known to cope with nitrogen starvation, were observed in both vegetation zones as well as Gaiellaceae and Sphingomonadaceae, bacteria associated with phosphorous limitations or metal pollution, respectively. (72,73). Thus, although a certain level of divergence was noticed in the chemical rhizosphere markers between plant species, similar ecological consequences on the soil microbiome were found. However, in the Prepuna, more correlations with growth-promoting bacteria were found. These growth-promotiong bacteria may be beneficial for annual species mostly found at low altitudes (*versus* perennials in Steppe), as they may promote faster development to escape periods of drought (36). In addition, some of the correlated bacterial families have been shown to relate to plant metabolism, which is supported by our analytical approach. For instance, coumarins and organic acids have been associated with Oxalobacteraceae, Comamonadaceae and pathogen taxa, respectively (7,32,74). Flavonoids were shown to mediate interactions with nitrogen fixers such as Oxalobacteraceae (75), a connection also observed in the present study. Finally, intriguing results arose from the analysis of soil samples collected in open areas. Richness and Shannon’s diversity were high in soil collected in the Steppe and metabolic markers from *J. frigida* rhizospheres could partially predict the elevation of samples from open areas, while *A. imbricata* rhizosphere markers failed. The abundance of rhizosphere chemicals from *J. frigida* was also correlated to significant changes in OTU abundances in open areas of the Steppe. Perennial Poaceae, such as *J. frigida*, can develop vegetative runners belowground, which may explain a diffusion of root chemicals. The implications of this diffusion and its relation with soil microbiome on Atacama dynamics needs further investigation, especially since an influence of plant-soil feedback on heterospecific plant successors has been shown (76).

Working with wild species in natural environments, particularly in an extreme environment such as the Atacama Desert, is very challenging. Moreover, these species are not easily grown or reproduced under laboratory conditions. However, field studies can provide key biological insights that are not achievable under laboratory conditions. This work provides unexpected insights into the response of rhizosphere chemistry to environmental constraints: direct (*i.e.* similar markers), as well as indirect (*i.e.* similar consequences on soil microbiome) convergence, could be observed between plant species. Rhizosphere chemistry predicted plant environment independently of the sampling year and, within a vegetation belt, chemical markers from one plant species could be used to predict the elevation level of other species. In addition, predictive metabolomics combined with OTU abundance analyses highlighted the diffusion of the rhizosphere markers of *J. frigida* and its implications for microbe communities. Rhizosphere chemistry and its influence on conspecific and heterospecific neighbours deserve greater attention to better predict ecosystem dynamics in the Atacama Desert and other extreme ecosystems.

## Acknowledgements

We are grateful to the indigenous community of Talabre for their authorisation to access the TLT transect. We also acknowledge the Genotoul bioinformatics platform Toulouse Occitanie (Bioinfo Genotoul, https://doi.org/10.15454/1.5572369328961167E12) for providing computing resources. This work was funded by the ANID-Millenuym Science Initiative Program-iBio ICN17_022; IEB (FB210006) and CGR ICN2021_044. We are also grateful to the Bordeaux Metabolome Facility (https://doi.org/10.15454/1.5572412770331912E12) and the MetaboHUB (ANR-11-INBS-0010) project for their support. CM also acknowledges the German Research Foundation (DFG, project MU 1829/29-1) for financial support.

## Author contributions

CL and RAG conceived the Atacama project. TD, RAG and PP conceived this analysis. Sampling was performed by RD, CL, FPD and RAG. TD and MCBS performed the LC-MS measurements. Statistical analyses on metabolomics data and OTU abundances were performed by TD, CL, CAN, SP, MG, CM, DR, RAG and PP. Metabolic markers and OTU abundances were linked by TD, CAN, MG and RAG. TD, RAG and PP wrote the manuscript with feedback from all co-authors.

## Data Availability

Data and metadata are included in supplemental tables and were deposited on MassIVE (MSV000093102, doi:10.25345/C5JM23S4X), which include raw spectra, non-normalised dataset, and feature metadata in the sections “raw”, “other” and “metadata”, respectively. Sample metadata are available in Tab. S1.

## Notes

### Competing Interest Statement

The authors have declared no competing interest.

## References

1. van Dam NM. How plants cope with biotic interactions. Plant Biology. 2009 Jan;11(1):1–5.

2. Kraiser T, Gras DE, Gutiérrez AG, González B, Gutiérrez RA. A holistic view of nitrogen acquisition in plants. Journal of Experimental Botany. 2011 Feb 1;62(4):1455–66.

3. Trivedi P, Leach JE, Tringe SG, Sa T, Singh BK. Plant–microbiome interactions: from community assembly to plant health. Nature Reviews Microbiology. 2020 Nov;18(11):607–21.

4. Berendsen RL, Pieterse CMJ, Bakker PAHM. The rhizosphere microbiome and plant health. Trends in Plant Science. 2012 Aug;17(8):478–86.

5. Oburger E, Schmidt H. New methods to unravel rhizosphere processes. Trends in Plant Science. 2016 Mar;21(3):243–55.

6. Bennett JA, Klironomos J. Mechanisms of plant–soil feedback: interactions among biotic and abiotic drivers. New Phytologist. 2019 Apr;222(1):91–6.

7. Yu P, He X, Baer M, Beirinckx S, Tian T, Moya YAT, et al. Plant flavones enrich rhizosphere *Oxalobacteraceae* to improve maize performance under nitrogen deprivation. Nature Plants. 2021 Apr;7(4):481–99.

8. Vismans G, van Bentum S, Spooren J, Song Y, Goossens P, Valls J, et al. Coumarin biosynthesis genes are required after foliar pathogen infection for the creation of a microbial soil-borne legacy that primes plants for SA-dependent defenses. Scientific Reports. 2022 Dec 28;12(1):22473.

9. Jones MD, Smith SE. Exploring functional definitions of mycorrhizas: are mycorrhizas always mutualisms? Canadian Journal of Botany. 2004 Aug 1;82(8):1089–109.

10. Gibert A, Tozer W, Westoby M. Plant performance response to eight different types of symbiosis. New Phytologist. 2019 Apr;222(1):526–42.

11. Hartmann M, Six J. Soil structure and microbiome functions in agroecosystems. Nature Reviews Earth & Environment. 2022 Nov 22;4(1):4–18.

12. Cosme M. Mycorrhizas drive the evolution of plant adaptation to drought. Communications Biology. 2023 Mar 30;6(1):346.

13. Hartmann A, Rothballer M, Schmid M. Lorenz Hiltner, a pioneer in rhizosphere microbial ecology and soil bacteriology research. Plant Soil. 2008 Nov;312(1–2):7–14.

14. Cotton TEA, Pétriacq P, Cameron DD, Meselmani MA, Schwarzenbacher R, Rolfe SA, et al. Metabolic regulation of the maize rhizobiome by benzoxazinoids. ISME Journal 2019 Jul;13(7):1647–58.

15. McLaughlin S, Zhalnina K, Kosina S, Northen TR, Sasse J. The core metabolome and root exudation dynamics of three phylogenetically distinct plant species. Nature Communications. 2023 Mar 24;14(1):1649.

16. Badri DV, Vivanco JM. Regulation and function of root exudates. Plant, Cell & Environment. 2009 Jun;32(6):666–81.

17. Fitzpatrick CR, Salas-González I, Conway JM, Finkel OM, Gilbert S, Russ D, et al. The plant microbiome: from ecology to reductionism and beyond. Annual Review of Microbiology. 2020 Sep 8;74(1):81–100.

18. Pétriacq P, Williams A, Cotton A, McFarlane AE, Rolfe SA, Ton J. Metabolite profiling of non-sterile rhizosphere soil. Plant Journal 2017 Oct;92(1):147–62.

19. Carvalhais LC, Dennis PG, Fedoseyenko D, Hajirezaei M, Borriss R, von Wirén N. Root exudation of sugars, amino acids, and organic acids by maize as affected by nitrogen, phosphorus, potassium, and iron deficiency. Z Pflanzenernähr Bodenk. 2011 Feb;174(1):3–11.

20. Frémont A, Sas E, Sarrazin M, Gonzalez E, Brisson J, Pitre FE, et al. Phytochelatin and coumarin enrichment in root exudates of arsenic-treated white lupin. Plant Cell & Environment. 2022 Mar;45(3):936–54.

21. Luginbuehl LH, Menard GN, Kurup S, Van Erp H, Radhakrishnan GV, Breakspear A, et al. Fatty acids in arbuscular mycorrhizal fungi are synthesized by the host plant. Science. 2017 Jun 16;356(6343):1175–8.

22. Sasse J, Martinoia E, Northen T. Feed Your Friends: Do plant exudates shape the root microbiome? Trends in Plant Science. 2018 Jan;23(1):25–41.

23. Macías FA, Marín D, Oliveros-Bastidas A, Castellano D, Simonet AM, Molinillo JMG. Structure−activity relationships (SAR) studies of benzoxazinones, their degradation products and analogues. Phytotoxicity on standard target species (STS). Journal of Agricultural and Food Chemistry. 2005 Feb 1;53(3):538–48.

24. Steinauer K, Thakur MP, Emilia Hannula S, Weinhold A, Uthe H, van Dam NM, et al. Root exudates and rhizosphere microbiomes jointly determine temporal shifts in plant-soil feedbacks. Plant Cell & Environment. 2023 Feb 27;pce.14570.

25. Schandry N, Becker C. Allelopathic Plants: Models for studying plant–interkingdom interactions. Trends in Plant Science. 2020 Feb;25(2):176–85.

26. Hawkins HJ, Cargill RIM, Van Nuland ME, Hagen SC, Field KJ, Sheldrake M, et al. Mycorrhizal mycelium as a global carbon pool. Current Biology. 2023 Jun;33(11):R560–73.

27. Delory BM, Delaplace P, Fauconnier ML, du Jardin P. Root-emitted volatile organic compounds: can they mediate belowground plant-plant interactions? Plant Soil. 2016 May;402(1–2):1–26.

28. Baldrian P, López-Mondéjar R, Kohout P. Forest microbiome and global change. Nature Reviews Microbiology. 2023 Mar 20;21-487–501).

29. Mandakovic D, Aguado-Norese C, García-Jiménez B, Hodar C, Maldonado JE, Gaete A, et al. Testing the stress gradient hypothesis in soil bacterial communities associated with vegetation belts in the Andean Atacama Desert. Environmental Microbiome. 2023 Mar 28;18(1):24.

30. Ruiz-Lozano JM, Aroca R, Zamarreño ÁM, Molina S, Andreo-Jiménez B, Porcel R, et al. Arbuscular mycorrhizal symbiosis induces strigolactone biosynthesis under drought and improves drought tolerance in lettuce and tomato: Drought and AM symbiosis induce strigolactones. Plant Cell & Environment. 2016 Feb;39(2):441–52.

31. Gargallo-Garriga A, Preece C, Sardans J, Oravec M, Urban O, Peñuelas J. Root exudate metabolomes change under drought and show limited capacity for recovery. Scientific Reports. 2018 Aug 23;8(1):12696.

32. Del Valle I, Webster TM, Cheng HY, Thies JE, Kessler A, Miller MK, et al. Soil organic matter attenuates the efficacy of flavonoid-based plant-microbe communication. Science Advances. 2020 Jan 31;6(5):eaax8254.

33. Branco S, Schauster A, Liao H, Ruytinx J. Mechanisms of stress tolerance and their effects on the ecology and evolution of mycorrhizal fungi. New Phytologist. 2022 Sep;235(6):2158–75.

34. Trivedi P, Batista BD, Bazany KE, Singh BK. Plant–microbiome interactions under a changing world: responses, consequences and perspectives. New Phytologist. 2022 Jun;234(6):1951–9.

35. Eshel G, Araus V, Undurraga S, Soto DC, Moraga C, Montecinos A, et al. Plant ecological genomics at the limits of life in the Atacama Desert. Proceedings of the National Academy of Sciences. 2021 Nov 16;118(46):e2101177118.

36. Díaz FP, Latorre C, Carrasco-Puga G, Wood JR, Wilmshurst JM, Soto DC, et al. Multiscale climate change impacts on plant diversity in the Atacama Desert. Global Change Biology. 2019 May 1;25(5):1733–45.

37. Luna E, Flandin A, Cassan C, Prigent S, Chevanne C, Kadiri CF, et al. Metabolomics to exploit the primed immune system of tomato fruit. Metabolites. 2020 Mar;10(3):96.

38. Dussarrat T, Prigent S, Latorre C, Bernillon S, Flandin A, Díaz FP, et al. Predictive metabolomics of multiple Atacama plant species unveils a core set of generic metabolites for extreme climate resilience. New Phytologist. 2022 Mar 15;nph.18095;234(1614–1628).

39. Tsugawa H, Cajka T, Kind T, Ma Y, Higgins B, Ikeda K, et al. MS-DIAL: data-independent MS/MS deconvolution for comprehensive metabolome analysis. Nature Methods. 2015 Jun;12(6):523–6.

40. Xia J, Sinelnikov IV, Han B, Wishart DS. MetaboAnalyst 3.0--making metabolomics more meaningful. Nucleic Acids Research. 2015 Jul 1;43(W1):W251–257.

41. R Core Team. R: a language and environment for statistical computing. Vienna, Austria: R Foundation for Statistical Computing; 2022.

42. Lê S, Josse J, Husson F. FactoMineR : An *R* package for multivariate analysis. Journal of Statistical Software. 2008; 25(1-18).

43. Petrén H, Köllner TG, Junker RR. Quantifying chemodiversity considering biochemical and structural properties of compounds with the R package CHEMODIV. New Phytologist. 2023 Mar;237(6):2478–92.

44. Harrel, Jr. F. E. Hmisc: Harrell Miscellaneous

45. Liland KH, Mevik BH, Wehrens R. pls: Partial least squares and principal component regression. 2022.

46. Mendiburu F de. agricolae: Statistical Procedures for Agricultural Research. 2020.

47. Wickham H. ggplot2: elegant graphics for data analysis. Springer-Verlag New York; 2016.

48. Kassambara A. ggpubr: ‘ggplot2’ Based publication ready plots. 2020.

49. Ruttkies C, Schymanski EL, Wolf S, Hollender J, Neumann S. MetFrag relaunched: incorporating strategies beyond in silico fragmentation. Journal of Cheminformatics. 2016 Dec;8(1):3.

50. Hastings J, Owen G, Dekker A, Ennis M, Kale N, Muthukrishnan V, et al. ChEBI in 2016: Improved services and an expanding collection of metabolites. Nucleic Acids Research. 2016 Jan 4;44(D1):D1214–9.

51. Horai H, Arita M, Kanaya S, Nihei Y, Ikeda T, Suwa K, et al. MassBank: a public repository for sharing mass spectral data for life sciences. Journal of Mass Spectrometry. 2010 Jul 7;45(7):703–14.

52. Sumner LW, Lei Z, Nikolau BJ, Saito K, Roessner U, Trengove R. Proposed quantitative and alphanumeric metabolite identification metrics. Metabolomics. 2014 Dec 1;10(6):1047–9.

53. Dussarrat T, Schweiger R, Ziaja D, Nguyen TTN, Krause L, Jakobs R, et al. Influences of chemotype and parental genotype on metabolic fingerprints of tansy plants uncovered by predictive metabolomics. Sci Rep. 2023 Jul 19;13(1):11645.

54. Djoumbou Feunang Y, Eisner R, Knox C, Chepelev L, Hastings J, Owen G, et al. ClassyFire: automated chemical classification with a comprehensive, computable taxonomy. Journal of Cheminformatics. 2016 Dec;8(1):61.

55. Douglas GM, Maffei VJ, Zaneveld JR, Yurgel SN, Brown JR, Taylor CM, et al. PICRUSt2 for prediction of metagenome functions. Nature Biotechnology. 2020 Jun;38(6):685–8.

56. Van Nuland ME, Bailey JK, Schweitzer JA. Divergent plant–soil feedbacks could alter future elevation ranges and ecosystem dynamics. Nature Ecology & Evolution. 2017 Apr 28;1(6):0150.

57. Chung YA. The temporal and spatial dimensions of plant–soil feedbacks. New Phytologist. 2023 Mar;237(6):2012–9.

58. Jansson JK, Hofmockel KS. Soil microbiomes and climate change. Nature Reviews Microbiology. 2020 Jan;18(1):35–46.

59. Kleine S, Müller C. Drought stress and leaf herbivory affect root terpenoid concentrations and growth of *Tanacetum vulgare*. Journal of Chemical Ecology. 2014 Oct;40(10):1115–25.

60. Izaguirre-Mayoral ML, Lazarovits G, Baral B. Ureide metabolism in plant-associated bacteria: purine plant-bacteria interactive scenarios under nitrogen deficiency. Plant Soil. 2018 Jul;428(1– 2):1–34.

61. Kaur H, Chowrasia S, Gaur VS, Mondal TK. Allantoin: emerging role in plant abiotic stress tolerance. Plant Molecular Biology Reporter. 2021 Sep;39(3):648–61.

62. Long M, Shou J, Wang J, Hu W, Hannan F, Mwamba TM, et al. Ursolic acid limits salt-induced oxidative damage by interfering with nitric oxide production and oxidative defense machinery in rice. Frontiers in Plant Science. 2020 Jun 24;11:697.

63. Hassan S, Mathesius U. The role of flavonoids in root-rhizosphere signalling: opportunities and challenges for improving plant-microbe interactions. Journal of Experimental Botany. 2012 May 1;63(9):3429–44.

64. Siebers M, Brands M, Wewer V, Duan Y, Hölzl G, Dörmann P. Lipids in plant–microbe interactions. Biochimica et Biophysica Acta (BBA) - Molecular and Cell Biology of Lipids. 2016 Sep;1861(9):1379–95.

65. Wandrag EM, Bates SE, Barrett LG, Catford JA, Thrall PH, Putten WH, et al. Phylogenetic signals and predictability in plant–soil feedbacks. New Phytologist. 2020 Nov;228(4):1440–9.

66. Walker TWN, Alexander JM, Allard P, Baines O, Baldy V, Bardgett RD, et al. Functional Traits 2.0: The power of the metabolome for ecology. Journal of Ecology. 2022 Jan;110(1):4–20.

67. Xu S, Wang J, Guo Z, He Z, Shi S. Genomic convergence in the adaptation to extreme environments. Plant Communications. 2020 Nov;1(6):100117.

68. Angulo V, Beriot N, Garcia-Hernandez E, Li E, Masteling R, Lau JA. Plant–microbe eco-evolutionary dynamics in a changing world. New Phytologist. 2022 Jun;234(6):1919–28.

69. Philippot L, Raaijmakers JM, Lemanceau P, Van Der Putten WH. Going back to the roots: the microbial ecology of the rhizosphere. Nature Reviews Microbiology. 2013 Nov;11(11):789–99.

70. Carrión VJ, Perez-Jaramillo J, Cordovez V, Tracanna V, De Hollander M, Ruiz-Buck D, et al. Pathogen-induced activation of disease-suppressive functions in the endophytic root microbiome. Science. 2019 Nov;366(6465):606–12.

71. Xu Y, Zheng C, Liang L, Yi Z, Xue S. Quantitative assessment of the potential for soil improvement by planting *Miscanthus* on saline-alkaline soil and the underlying microbial mechanism. GCB Bioenergy. 2021 Jul;13(7):1191–205.

72. Wang W, Wang J, Wang Q, Bermudez RS, Yu S, Bu P, et al. Effects of plantation type and soil depth on microbial community structure and nutrient cycling function. Frontiers in Microbiology. 2022 May 31;13:846468.

73. Sultana R, Islam SMN, Sultana T. Arsenic and other heavy metals resistant bacteria in rice ecosystem: potential role in promoting plant growth and tolerance to heavy metal stress. Environmental Technology & Innovation. 2023 Aug;31:103160.

74. Wen T, Yuan J, He X, Lin Y, Huang Q, Shen Q. Enrichment of beneficial cucumber rhizosphere microbes mediated by organic acid secretion. Horticulture Research. 2020 Dec;7(1):154.

75. Hong Y, Zhou Q, Hao Y, Huang AC. Crafting the plant root metabolome for improved microbe-assisted stress resilience. New Phytologist. 2022 Jun;234(6):1945–50.

76. Wilschut RA, Hume BCC, Mamonova E, Van Kleunen M. Plant–soil feedback effects on conspecific and heterospecific successors of annual and perennial Central European grassland plants are correlated. Nat Plants. 2023 Jun 9(7):1057–1066.

