## Supplementary material for "Convergent and divergent responses of the rhizosphere chemistry and bacterial communities to a stress gradient in the Atacama Desert": Figure S1

### Slide 1
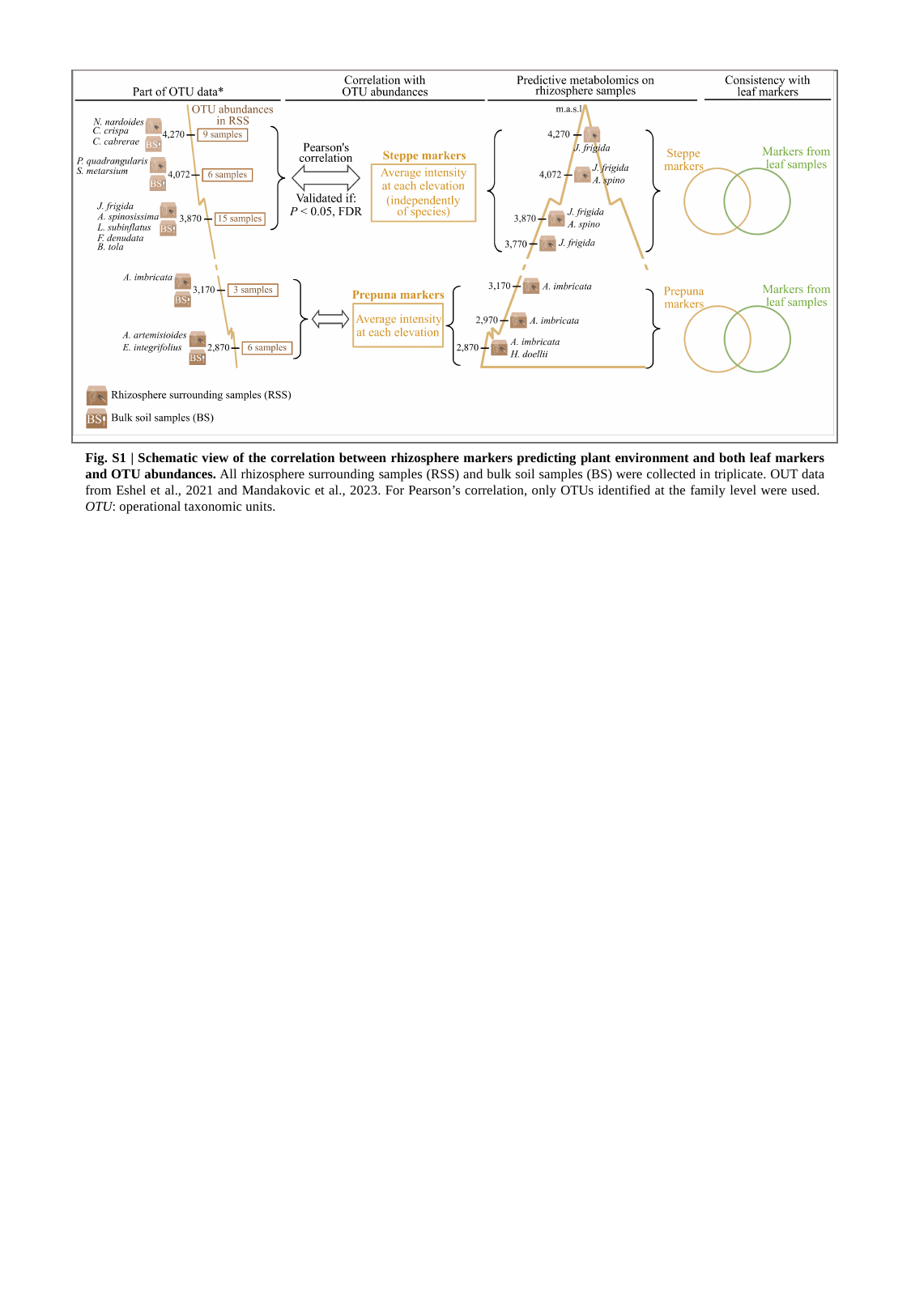

Fig. S1 | Schematic view of the correlation between rhizosphere markers predicting plant environment and both leaf markers and OTU abundances. All rhizosphere surrounding samples (RSS) and bulk soil samples (BS) were collected in triplicate. OUT data from Eshel et al., 2021 and Mandakovic et al., 2023. For Pearson’s correlation, only OTUs identified at the family level were used. OTU: operational taxonomic units.
