## Supplementary material for "Convergent and divergent responses of the rhizosphere chemistry and bacterial communities to a stress gradient in the Atacama Desert": Figure S2

### Slide 1
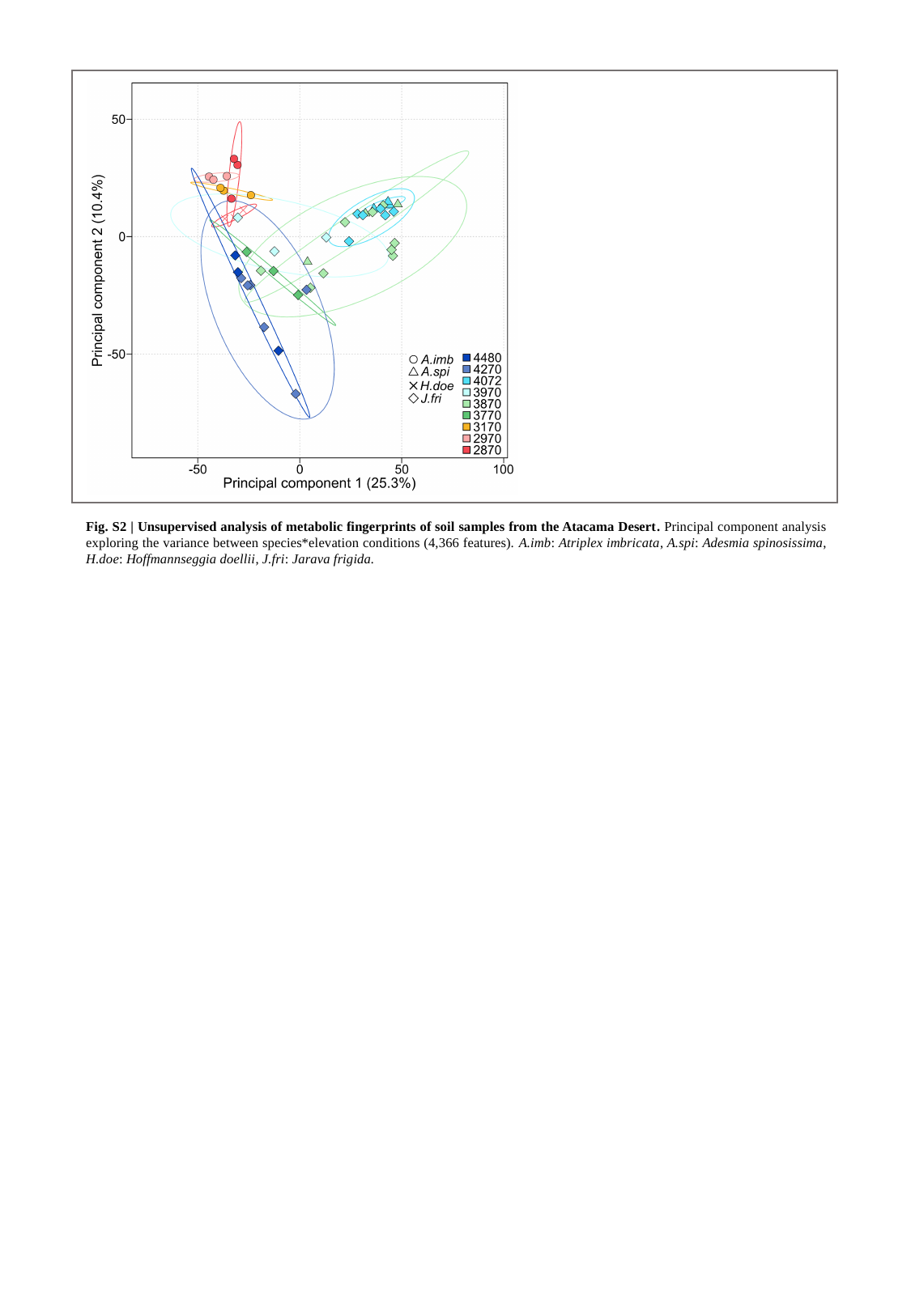

Fig. S2 | Unsupervised analysis of metabolic fingerprints of soil samples from the Atacama Desert. Principal component analysis exploring the variance between species*elevation conditions (4,366 features). A.imb: Atriplex imbricata, A.spi: Adesmia spinosissima, H.doe: Hoffmannseggia doellii, J.fri: Jarava frigida.
