## Supplementary material for "Convergent and divergent responses of the rhizosphere chemistry and bacterial communities to a stress gradient in the Atacama Desert": Figure S3

### Slide 1
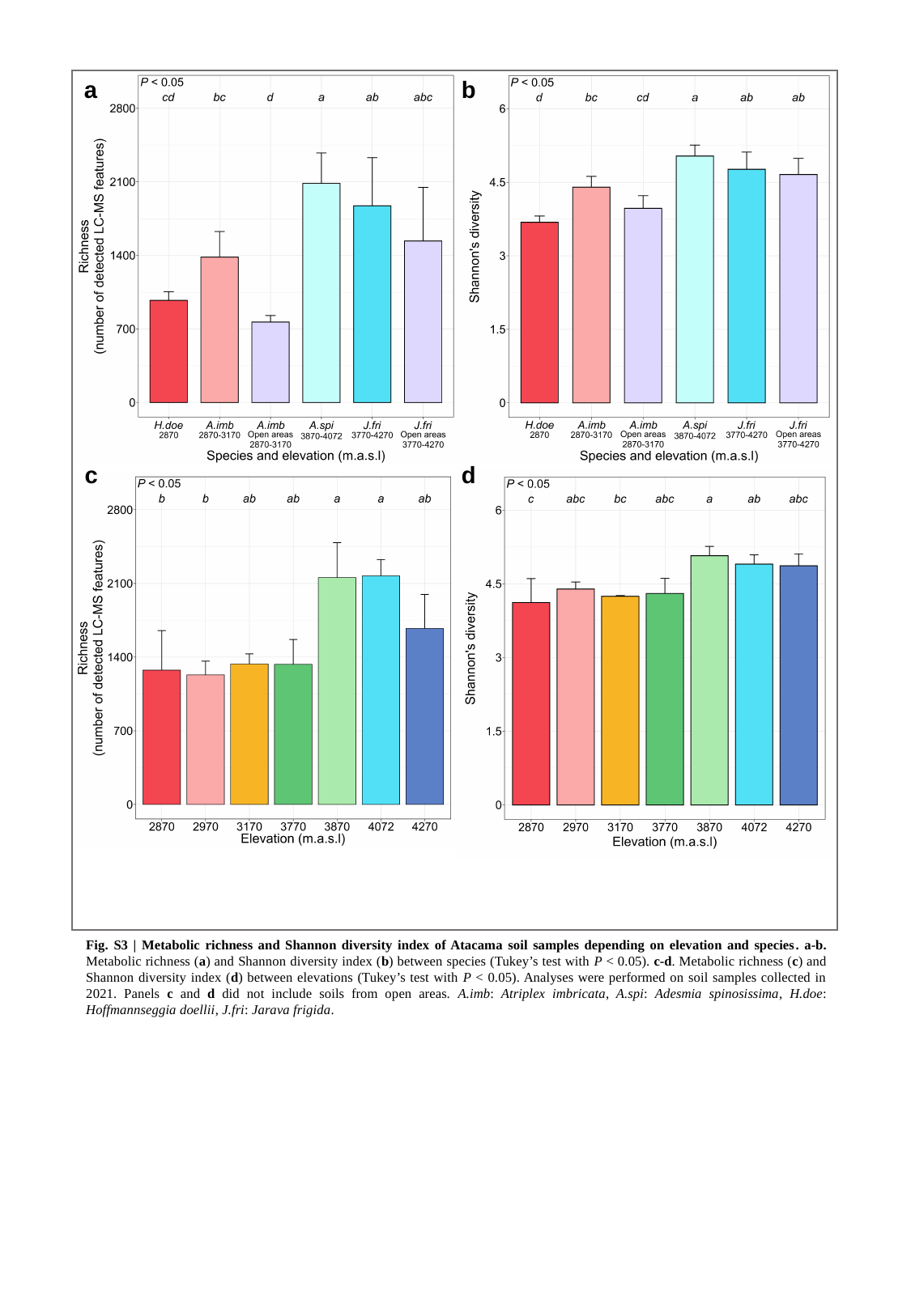

a
b
c
d
Fig. S3 | Metabolic richness and Shannon diversity index of Atacama soil samples depending on elevation and species. a-b. Metabolic richness (a) and Shannon diversity index (b) between species (Tukey’s test with P < 0.05). c-d. Metabolic richness (c) and Shannon diversity index (d) between elevations (Tukey’s test with P < 0.05). Analyses were performed on soil samples collected in 2021. Panels c and d did not include soils from open areas. A.imb: Atriplex imbricata, A.spi: Adesmia spinosissima, H.doe: Hoffmannseggia doellii, J.fri: Jarava frigida.
