## Supplementary material for "Convergent and divergent responses of the rhizosphere chemistry and bacterial communities to a stress gradient in the Atacama Desert": Figure S4

### Slide 1
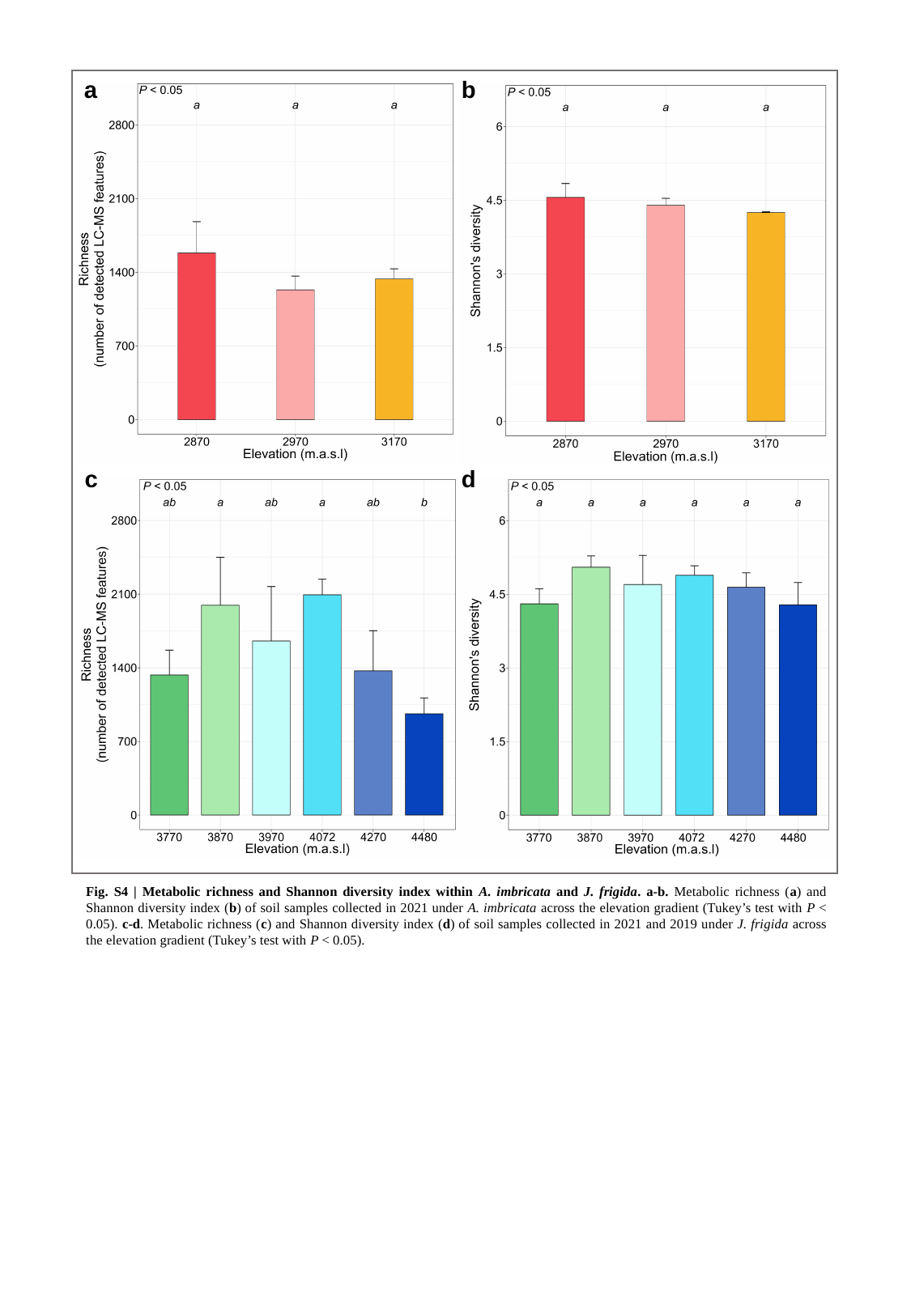

a
b
c
d
Fig. S4 | Metabolic richness and Shannon diversity index within A. imbricata and J. frigida. a-b. Metabolic richness (a) and Shannon diversity index (b) of soil samples collected in 2021 under A. imbricata across the elevation gradient (Tukey’s test with P < 0.05). c-d. Metabolic richness (c) and Shannon diversity index (d) of soil samples collected in 2021 and 2019 under J. frigida across the elevation gradient (Tukey’s test with P < 0.05).
