## Supplementary material for "Convergent and divergent responses of the rhizosphere chemistry and bacterial communities to a stress gradient in the Atacama Desert": Figure S5

### Slide 1
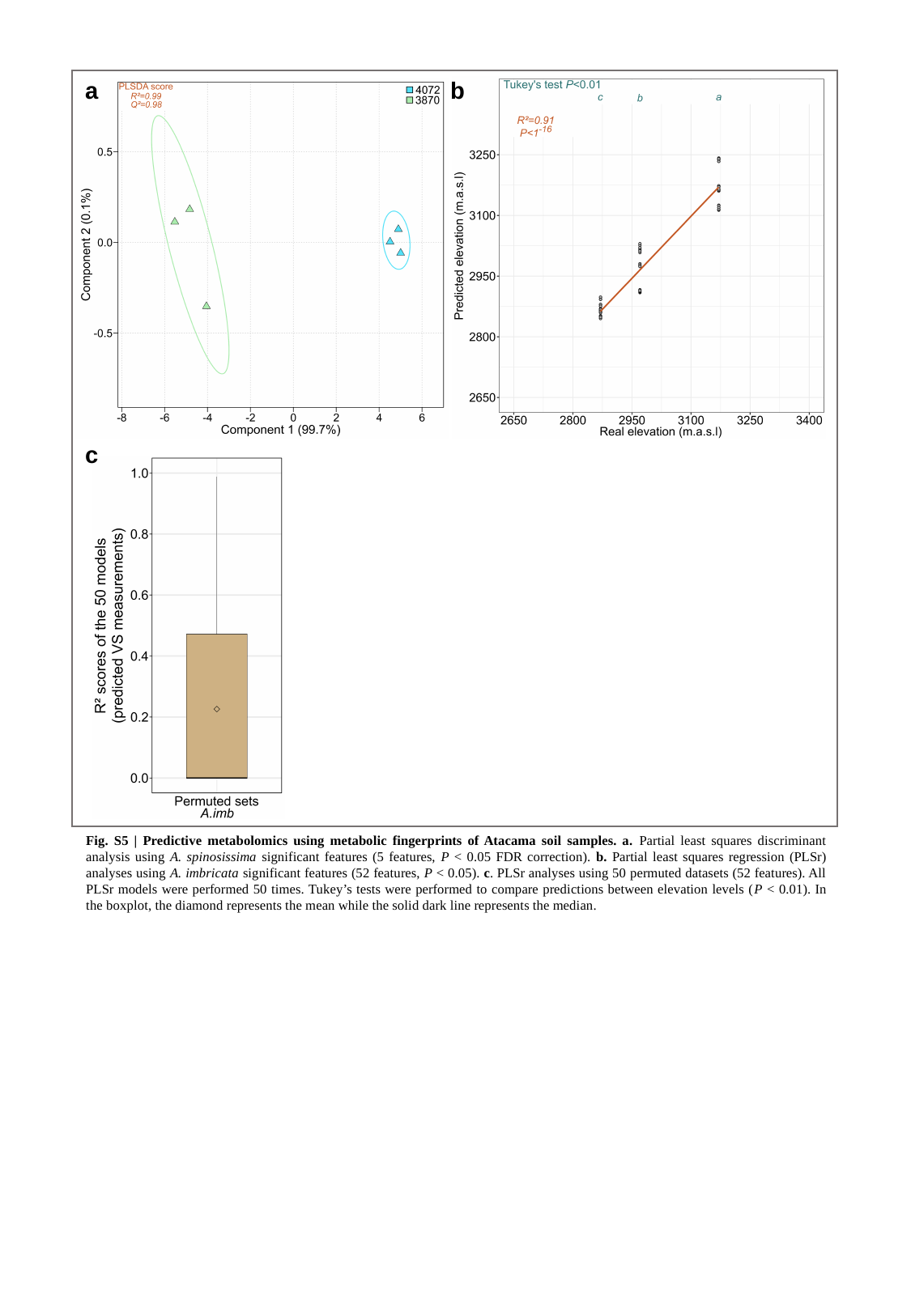

a
b
c
Fig. S5 | Predictive metabolomics using metabolic fingerprints of Atacama soil samples. a. Partial least squares discriminant analysis using A. spinosissima significant features (5 features, P < 0.05 FDR correction). b. Partial least squares regression (PLSr) analyses using A. imbricata significant features (52 features, P < 0.05). c. PLSr analyses using 50 permuted datasets (52 features). All PLSr models were performed 50 times. Tukey’s tests were performed to compare predictions between elevation levels (P < 0.01). In the boxplot, the diamond represents the mean while the solid dark line represents the median.
