## Supplementary material for "Convergent and divergent responses of the rhizosphere chemistry and bacterial communities to a stress gradient in the Atacama Desert": Figure S6

### Slide 1
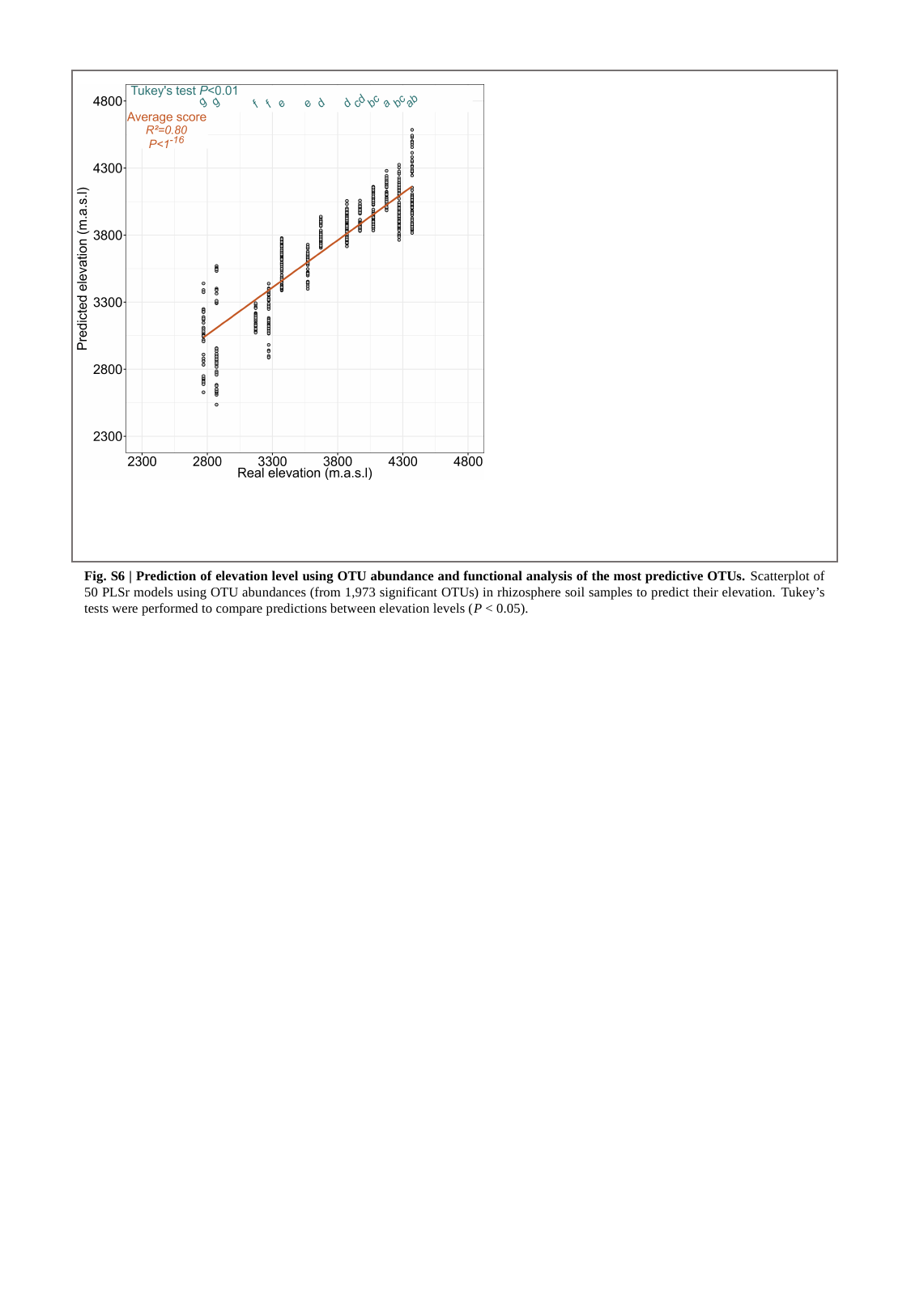

Fig. S6 | Prediction of elevation level using OTU abundance and functional analysis of the most predictive OTUs. Scatterplot of 50 PLSr models using OTU abundances (from 1,973 significant OTUs) in rhizosphere soil samples to predict their elevation. Tukey’s tests were performed to compare predictions between elevation levels (P < 0.05).
