## Supplementary material for "Convergent and divergent responses of the rhizosphere chemistry and bacterial communities to a stress gradient in the Atacama Desert": Figure S7

### Slide 1
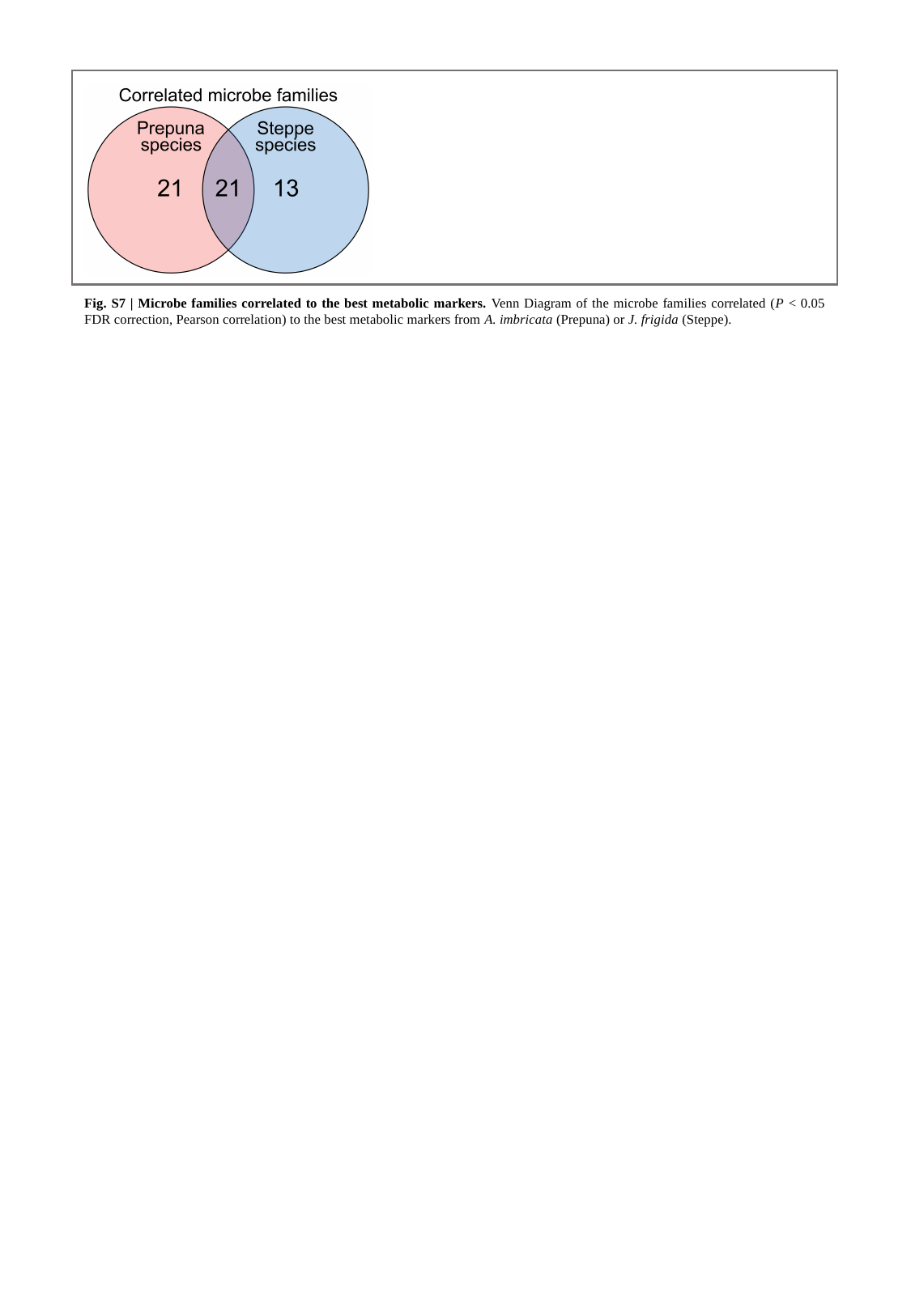

Fig. S7 | Microbe families correlated to the best metabolic markers. Venn Diagram of the microbe families correlated (P < 0.05 FDR correction, Pearson correlation) to the best metabolic markers from A. imbricata (Prepuna) or J. frigida (Steppe).
