## Supplementary material for "Convergent and divergent responses of the rhizosphere chemistry and bacterial communities to a stress gradient in the Atacama Desert": Figure S8

### Slide 1
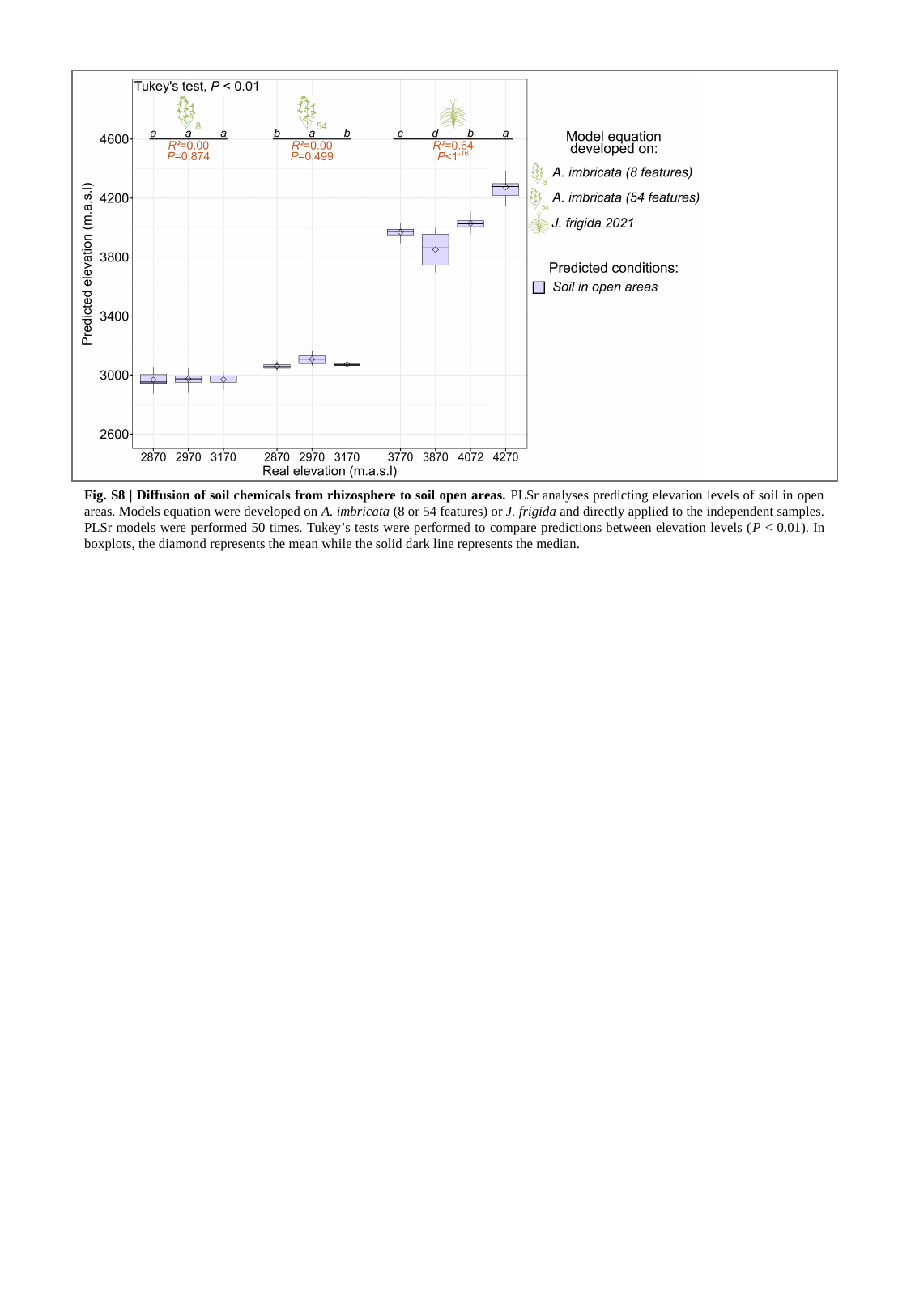

Fig. S8 | Diffusion of soil chemicals from rhizosphere to soil open areas. PLSr analyses predicting elevation levels of soil in open areas. Models equation were developed on A. imbricata (8 or 54 features) or J. frigida and directly applied to the independent samples. PLSr models were performed 50 times. Tukey’s tests were performed to compare predictions between elevation levels (P < 0.01). In boxplots, the diamond represents the mean while the solid dark line represents the median.
